# Mapping of Whole-Brain Resting-State Networks with Half-Millimetre Resolution

**DOI:** 10.1101/2021.03.09.434629

**Authors:** Seong Dae Yun, Patricia Pais-Roldán, Nicola Palomero-Gallagher, N. Jon Shah

**Affiliations:** Institute of Neuroscience and Medicine 4, Medical Imaging Physics, Forschungszentrum Jülich, Germany; Institute of Neuroscience and Medicine 1, Structural and Functional Organisation of the Brain, Forschungszentrum Jülich, Germany; C. & O. Vogt Institute for Brain Research, Heinrich-Heine-University, 40225 Düsseldorf, Germany; Department of Psychiatry, Psychotherapy and Psychosomatics, Medical Faculty, RWTH Aachen, Aachen, Germany; Institute of Neuroscience and Medicine 11, Molecular Neuroscience and Neuroimaging, JARA, Forschungszentrum Jülich, Germany; JARA - BRAIN - Translational Medicine, Aachen, Germany; Department of Neurology, RWTH Aachen University, Aachen, Germany

## Abstract

Resting-state fMRI has been used in numerous studies to map networks in the brain that employ spatially disparate regions. However, attempts to map networks with high spatial resolution have been hampered by conflicting technical demands and associated problems. Results from recent fMRI studies have shown that spatial resolution remains around 0.7 × 0.7 × 0.7 mm^3^, with only partial brain coverage. This work presents a novel fMRI method, TR-external EPI with keyhole (TR-external EPIK), which can provide a nominal spatial resolution of 0.51 × 0.51 × 1.00 mm^3^ (0.26 mm^3^ voxel) with whole-brain coverage. TR-external EPIK enabled the identification of various resting-state networks distributed throughout the brain from a single fMRI session, with mapping fidelity onto the grey matter at 7T. The high-resolution functional image further revealed mesoscale anatomical structures, such as small cerebral vessels and the internal granular layer of the cortex within the postcentral gyrus.

## 1. Introduction

Since the first demonstration of blood-oxygenated-level-dependent (BOLD) contrast using MRI [1], fMRI, as it has become known, has been widely used to explore brain function under resting-state, or evoked by a stimulus-driven paradigm. Task-evoked fMRI detects functional signals involved with a specific given task, which reflect only a small fraction of the brain’s overall activity [2]. In contrast, resting-state fMRI focuses on spontaneous neuronal activity fluctuating at very low frequencies (< 0.1 Hz), by which correlated brain areas in disparate regions, termed ‘resting-state networks’, can be identified [2,3]. Thus, resting-state fMRI has been employed by many groups for the investigation of overall brain function and its underlying connectivity [2]. In addition, the task-free acquisition of restingstate fMRI allows various neurological issues to be studied in patients (e.g. cognitive dysfunction, psychiatric disorders, consciousness, etc.) or young children (e.g. neonates, infants, etc.), who either have difficulties or simply cannot comply with designed paradigms [4–6].

For the identification of correlated brain areas in resting-state fMRI, whole-brain coverage is demanded as one of the first requirements. Results from several clinical applications have shown that consciousness impairment and neuropsychiatric disorders such as in schizophrenia or epilepsy patients are linked to the alteration of several brain networks that span multiple cortical areas [4,5,7]. This fact indicates a need for whole-brain imaging to detect resting-state interactions of potential diagnostic/prognostic value. For the analysis of brain connectivity across distant brain regions, a model-free (i.e. data-driven) approach, known as independent component analysis (ICA), is commonly used to effectively extract the spatially and temporally independent functional components from the whole brain [2,3].

Recent advances in fMRI techniques enable the depiction of neuronal activation with a submillimetre voxel size. Previously, several fMRI studies have employed high-resolution imaging techniques to measure the fMRI signal with a cortical depth-dependence (referred to as “laminar fMRI”, despite the absence of any direct correlation between the sampled depths and the histologically-defined cortical laminae) [8,9]. However, most of these high-resolution laminar fMRI investigations of activation profiles through the cortex have only targeted functional activation evoked by a task paradigm in a particular brain region [10–14], leaving the laminar dynamics underlying the resting-state largely unexplored. This is mainly due to the technical limitations of the current fMRI techniques, which are unable to offer submillimetre voxel size with whole-brain coverage, hindering the acquisition of wholebrain laminar fMRI data.

There have been numerous attempts to improve the spatiotemporal resolution of fMRI, including the use of a reduced FOV [15–17] or the use of non-Cartesian and random sampling trajectories [18–20]. Although a reduced FOV can usually achieve a higher spatial resolution than full-FOV schemes, the restricted FOV is a critical limiting factor in investigating whole-brain functionality. The non-Cartesian approaches improve spatial resolution by employing a more efficient k-space sampling scheme, such as radial or spiral trajectories [18,19]; however, they typically suffer from off-resonance artefacts, rendering a robust localisation of functional activity difficult without the application of an additional correction method [18]. Although the random sampling method also offers more robust image reconstruction, leading to a highly accelerated acquisition, its complex image reconstruction can introduce distortions to functional time-series data, resulting in the reduction of BOLD amplitudes [20].

For the reasons described above, full-FOV Cartesian sampling methods are still widely used in submillimetre-resolution fMRI studies, and of these, echo-planar imaging (EPI) is the most commonly implemented method [21–25]. However, the spatial resolution achieved in several recent works has remained around 0.7 × 0.7 × 0.7 mm^3^, and most methods are unable to provide whole-brain coverage.

This work presents a novel fMRI methodology, providing a half-millimetre in-plane resolution with whole-brain coverage. The imaging method was developed based on the combination of TR-external EPI phase correction [26,27] with EPI with keyhole (EPIK) [28–39], both of which have been shown to be effective in improving the spatial resolution and brain coverage while maintaining comparable BOLD detection performance when compared to a standard EPI method [27, 32,34–37]. The developed imaging method is termed “TR-external EPIK” [38,39].

This work focuses on the evaluation of a half-millimetre protocol implemented using TR-external EPIK at 7T. A qualitative inspection of the achieved spatial resolution was performed by identifying mesoscale anatomical structures in reconstructed images. The protocol was employed in resting-state fMRI to demonstrate high-resolution mapping of activated voxels in a number of resting-state networks distributed widely over the brain. Furthermore, resting-state fMRI data were acquired from a group of subjects to verify the robust detection of functional signals using TR-external EPIK. The imaging performance of TR-external EPIK was evaluated by comparison with the imaging parameters of previously published submillimetre-resolution fMRI studies.

## 2. Results

### 2.1. Functional scan with a half-millimetre in-plane resolution

An fMRI protocol yielding a 0.26 mm^3^ voxel (in-plane: 0.51 × 0.51 mm^2^) was implemented using TR-external EPIK. The protocol was configured to have a FOV (210 × 210 mm^2^) sufficient to cover the entire brain sliced in the axial direction while providing brain coverage of 108 mm in the dorso-ventral direction (108 slices; 1 mm thickness with no slice-gap). A more detailed description of the imaging protocol can be found in the Materials and Methods and Supporting Information (S1 Fig and S1 Table). Fig 1 shows images reconstructed from TR-external EPIK. Here, the surface of the grey matter (GM) and the white matter (WM) of the brain were extracted from all the acquired slices and rendered in 3D (Fig 1a), effectively demonstrating the near whole-brain coverage provided by TR-external EPIK.

**Fig 1.**
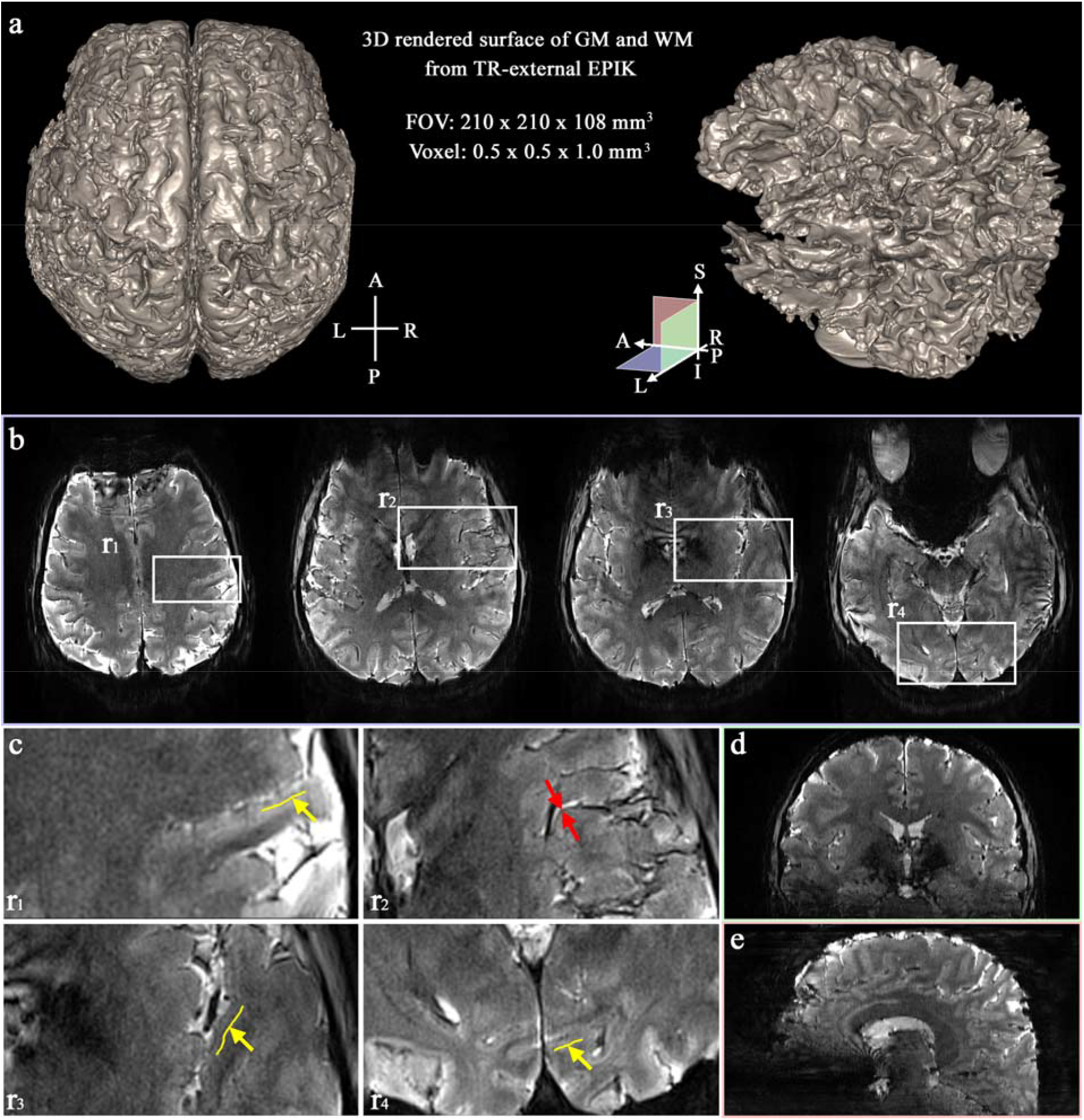
Reconstructed images from the half-millimetre (0.51 × 0.51 mm^2^) protocol. (**a**) 3D rendered surface of the GM (left) and WM (right) obtained from TR-external EPIK, (**b**) four representative slices, selected from a set of 108 axial slices, (**c**) enlarged depiction of the ROIs marked by the white rectangles in **b** (r_1_ ~ r_4_), where the following mesoscale anatomical structures can be observed: small cerebral vessels (r_2_) or the internal granular layer of the cortex within the postcentral gyrus (r_1_), the Heschl gyrus (r_3_) and the calcarine sulcus (r_4_) and (**d, e**) resliced and reconstructed coronal and sagittal views of the 108 axial slices demonstrating the extensive brain coverage provided by TR-external EPIK. Note that cortical ribbons are clearly seen even in the resliced images.

For more detailed visual inspection, sectional slices (axial, coronal and sagittal) were taken at their representative locations, as shown in Figs 1b, d and e. Fig 1b shows four of the 108 axial slices; the entire reconstructed slices can be found in S2 Fig. Here, specific brain regions were chosen (marked by white rectangles: r_1_~ r_4_) in which the following mesoscale anatomical structures can be observed (Fig 1c): small cerebral vessels (red arrows in r_2_) or the internal granular layer of the cortex (yellow arrows) located on the anterior wall of the postcentral gyrus (r_1_), on the Heschl gyrus (r_3_) or within the calcarine sulcus (r_4_). The complete extent of the brain covered by TR-external EPIK can be verified from the coronal and sagittal images (Figs 1d and e). It is important to note that the cortical ribbon can even be seen clearly in these resliced images. In addition, these depictions also show that all slices were reconstructed without any significant inter-slice artefacts that can sometimes result from the multi-band reconstruction [40]. The signal behaviour along the direction in which multi-band acceleration was applied (i.e. superior to inferior direction) is very continuous and does not depict any abrupt signal changes.

### 2.2. Mapping of Resting-state Networks

The above half-millimetre protocol was applied to record resting-state fMRI. Eighteen healthy volunteers participated in this study, and resting-state networks of each subject were obtained using independent component analysis (ICA). Fig 2 shows the results obtained from a representative subject, with Fig 2a showing the results for six resting-state fMRI networks (i.e. dorsal-DMN, ventral-DMN, visual, sensorimotor (LH), sensorimotor (RH) and fronto-parietal) selected from the independent components. The activated voxels are overlaid on the 3D rendered outer cortical surface, generated by segmenting the mean image of the re-aligned functional scans, in which the above resting-state networks in distant brain regions are clearly seen. Fig 2b shows the results presented in three different sectional views (axial, coronal and sagittal), where the activation pattern of each resting-state network is most clearly observed. Fig 2 clearly demonstrates that the extensive brain coverage provided by TR-external EPIK allowed the six resting-state networks to be simultaneously determined from a single fMRI session. In addition, the half-millimetre protocol (0.26 mm^3^ voxels) enabled the identification of functional voxels very locally along the cortical ribbon.

**Fig 2.**
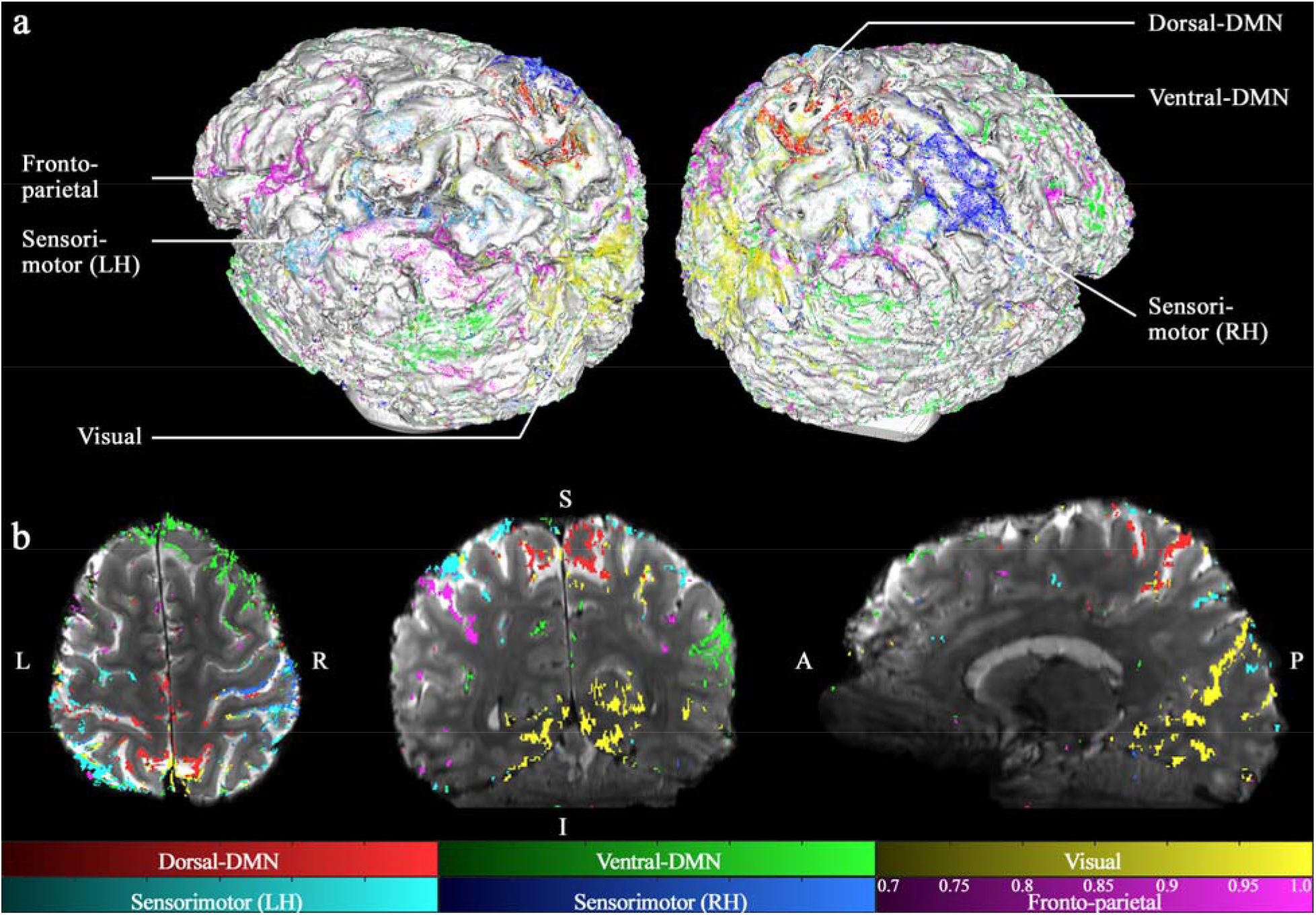
Whole-brain resting-state networks. The activated voxels were obtained with a statistical threshold (probability ≥ 0.7) and, for illustrative purpose, clusters containing less than three voxels were excluded from the visualisation. Resting-state results are shown for the following six selected networks: dorsal-DMN, ventral-DMN, visual, sensorimotor (LH), sensorimotor (RH) and fronto-parietal; here, DMN, LH and RH denote default mode network, left hemisphere and right hemisphere, respectively. The activated voxels are shown, overlaid on (**a**) a 3D rendered surface of the mean image of the re-aligned functional scans and (**b**) three sectional slices (axial, coronal and sagittal). This figure demonstrates the performance of TR-external EPIK in terms of brain coverage (108 mm in superior-inferior direction) and the localisation of the activated voxels on the cortical ribbons (nominal spatial resolution: 0.51 × 0.51 mm^2^).

For a more detailed examination, each resting-state network is displayed separately on its representative axial slice (Fig 3a); the results are also directly overlaid on the mean image of the realigned functional scans where non-brain tissues were eliminated using the standard brain extraction procedure in FSL (BET). Here, for the three networks shown in the right column of Fig 3a (i.e. dorsal-DMN, sensorimotor for the right hemisphere and visual), an ROI was selected from each network (marked with a green rectangle) and is displayed in a magnified view in Fig 3b. The size of the selected ROI is 40 × 40 voxels, which is approximately 20 × 20 mm^2^. This figure exhibits the localisation of the identified functional voxels along the cortical regions more clearly. Here, in order to aid the identification of cortical areas, the GM regions obtained from the standard segmentation routine in FSL are overlaid on the same figure, shown in blue. The localisation of functional voxels in the GM was further verified by plotting a line profile. As shown in Fig 3b, a line, starting from ‘P_1_’ and ending at ‘P_2_’ and crossing the cortical ribbon perpendicularly, was manually defined from each enlarged depiction, and the network-specific probability profile along this line was examined (the solid black line in Fig 3c). In Fig 3c, the signal intensity of the background image is also plotted (the black dotted line in Fig 3c) to serve as a reference for the T_2_/T_2_* along the underlying tissue. Here, signal intensity changes in the background image are observed, which gradually increase from the WM to the cerebrospinal fluid (CSF) region (i.e. T_2_/T_2_* contrast). The GM region obtained from the segmentation is also delineated here with a blue dashed line. For all three networks shown in Figs 3b and c, it can be seen that the BOLD activation is almost confined to the GM regions. In Fig 3b, it can be observed that some of the activated voxels are partly in the CSF region, which is in agreement with the known sensitivity of BOLD to the large pial vessels and is typically seen in gradient-echo-based sequences [41]. In this work, the effect caused by the susceptibility differences arising from pial vessels was reduced by applying a correction method for phase-related venous contributions [41,42].

**Fig 3.**
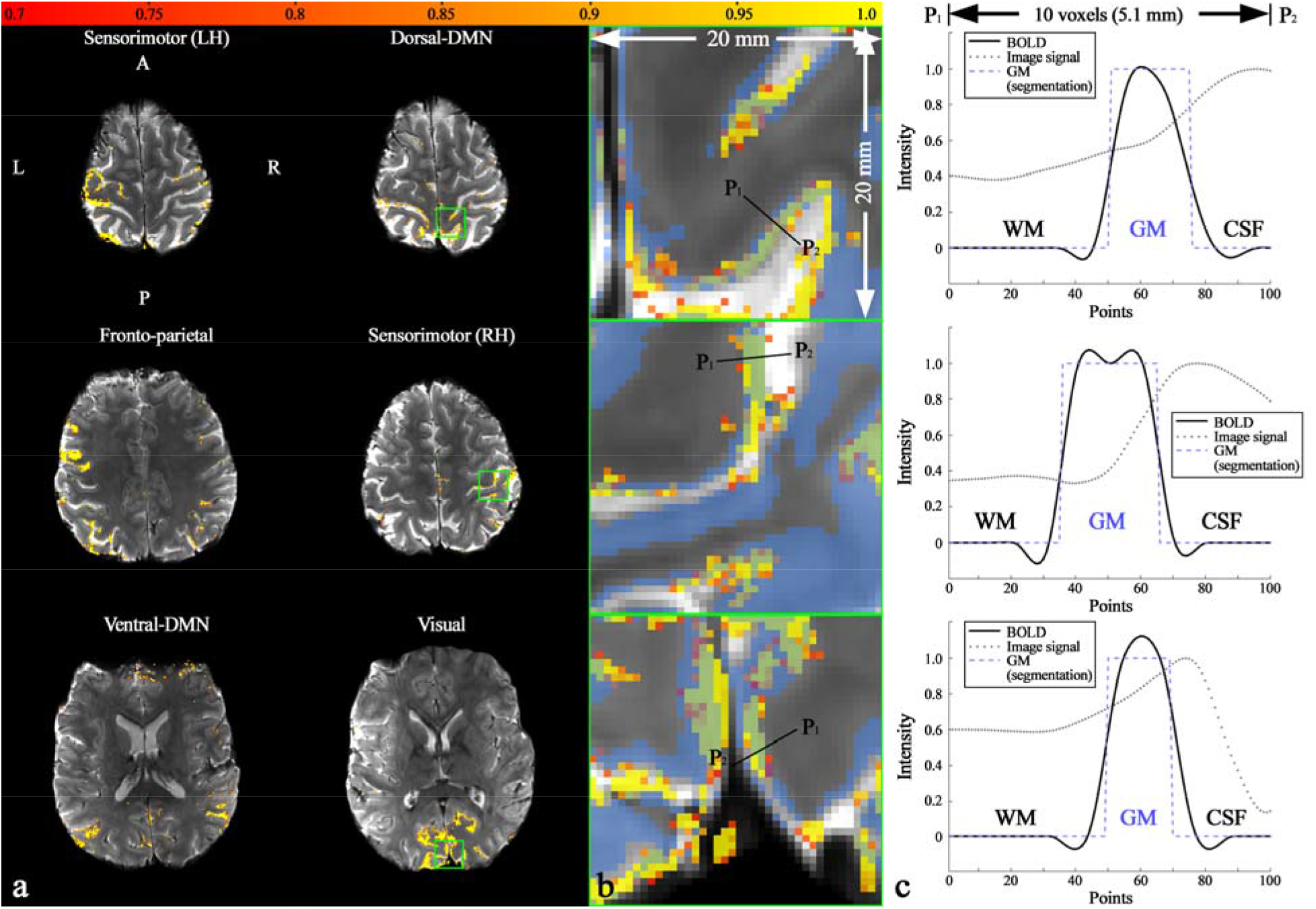
Results of the resting-state networks and line profiles. (**a**) Each resting-state network overlaid on its representative slice location, (**b**) enlarged depiction of the ROIs, marked by the green rectangles in the right column of Fig a (i.e. dorsal-DMN, sensorimotor (RH) and visual) and (**c**) line profile of the network-specific probability at the black line, denoted with the points P1 (start) and P2 (end). The length of the examined line is 10 voxels (5.1 mm), and 100 points were sampled along its length. The significance of the functional activation is coded with a red-to-yellow scale, as depicted on the top of the figure. To aid the identification of cortical regions, the GM obtained from segmentation is delineated in both Figs b and c in blue. It can be seen that the behaviour of BOLD signals is in line with the extent of the grey matter regions.

### 2.3. Resting-state networks from a group of subjects: reproducibility

Fig 4 summarises the resting-state networks obtained from 13 of the 18 subjects for six well-known functional networks: auditory, fronto-parietal, sensorimotor, visual, dorsal-DMN and ventral-DMN. The plot shows that although the detection pattern was different between subjects, the resting-state networks were reliably detected for all 13 subjects, which demonstrates the robust performance of the half-millimetre protocol in the detection of resting-state functional signals.

**Fig 4.**
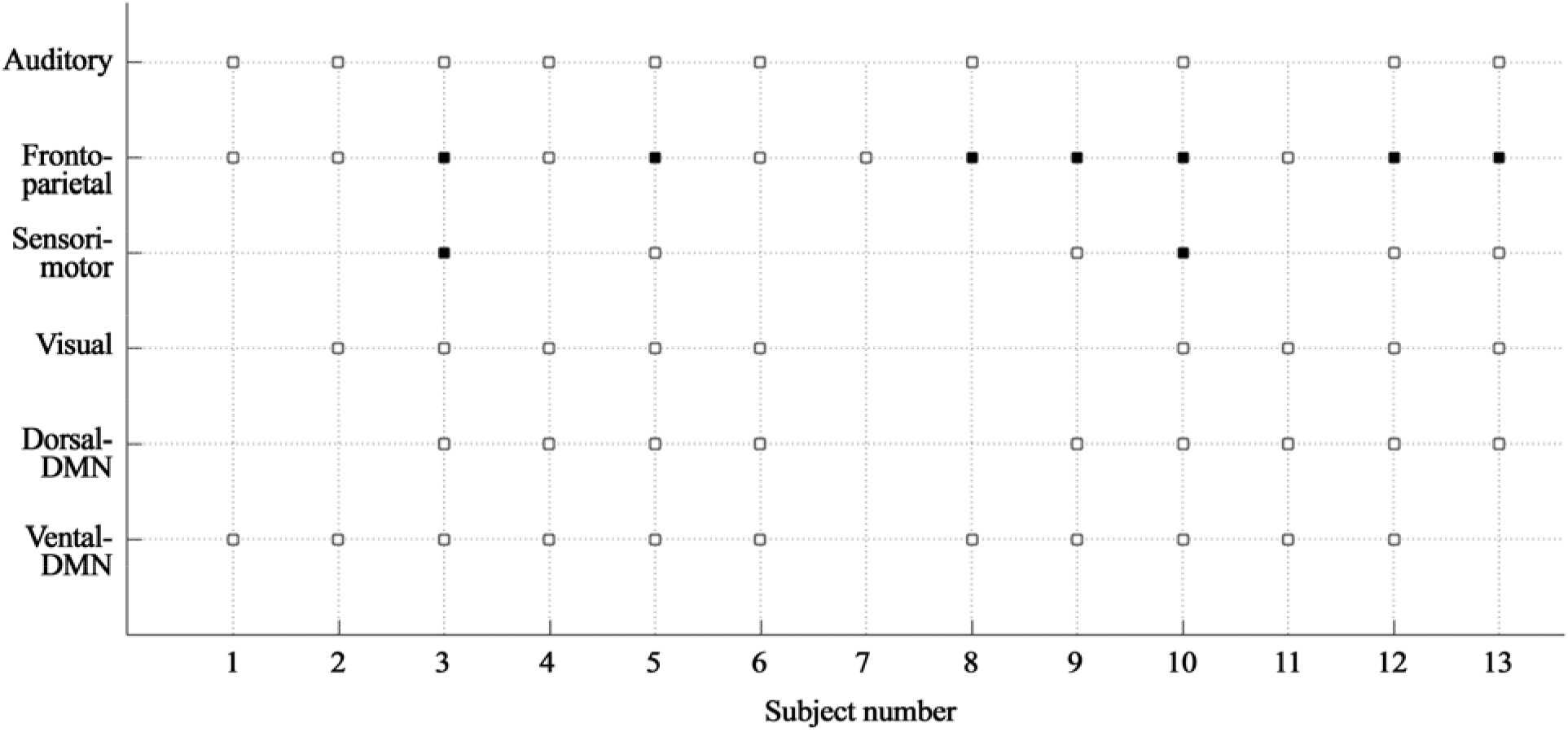
A grid plot of resting-state networks with respect to the 13 subjects. For six well-known functional networks (auditory, fronto-parietal, sensorimotor, visual, dorsal-DMN and ventral-DMN), the detected resting-state networks in each subject are marked with a square. The solid squares in the somatosensory and the parietal networks present the case when the activation was found in both hemispheres (i.e. bi-hemispheric activation); uni-hemispheric activation (either left or right) is depicted by the open squares. This figure depicts that the resting-state networks were reliably detected through all 13 subjects.

Fig 5 shows individually displayed, activated voxels, at their respective slice location for five of the subjects showing the same resting-state network (e.g., subjects 3, 5, 6, 8 and 12 for the auditory network). The analysis was performed for each of the following resting-state networks: auditory, frontoparietal, sensorimotor, visual, dorsal-DMN and ventral-DMN. The figure reveals that the identified networks had a similar activation pattern across different subjects, which demonstrates the reproducibility of mapping resting-state networks using TR-external EPIK. In addition, the activated voxels from each subject are also shown to be localised along the cortical ribbon.

**Fig 5.**
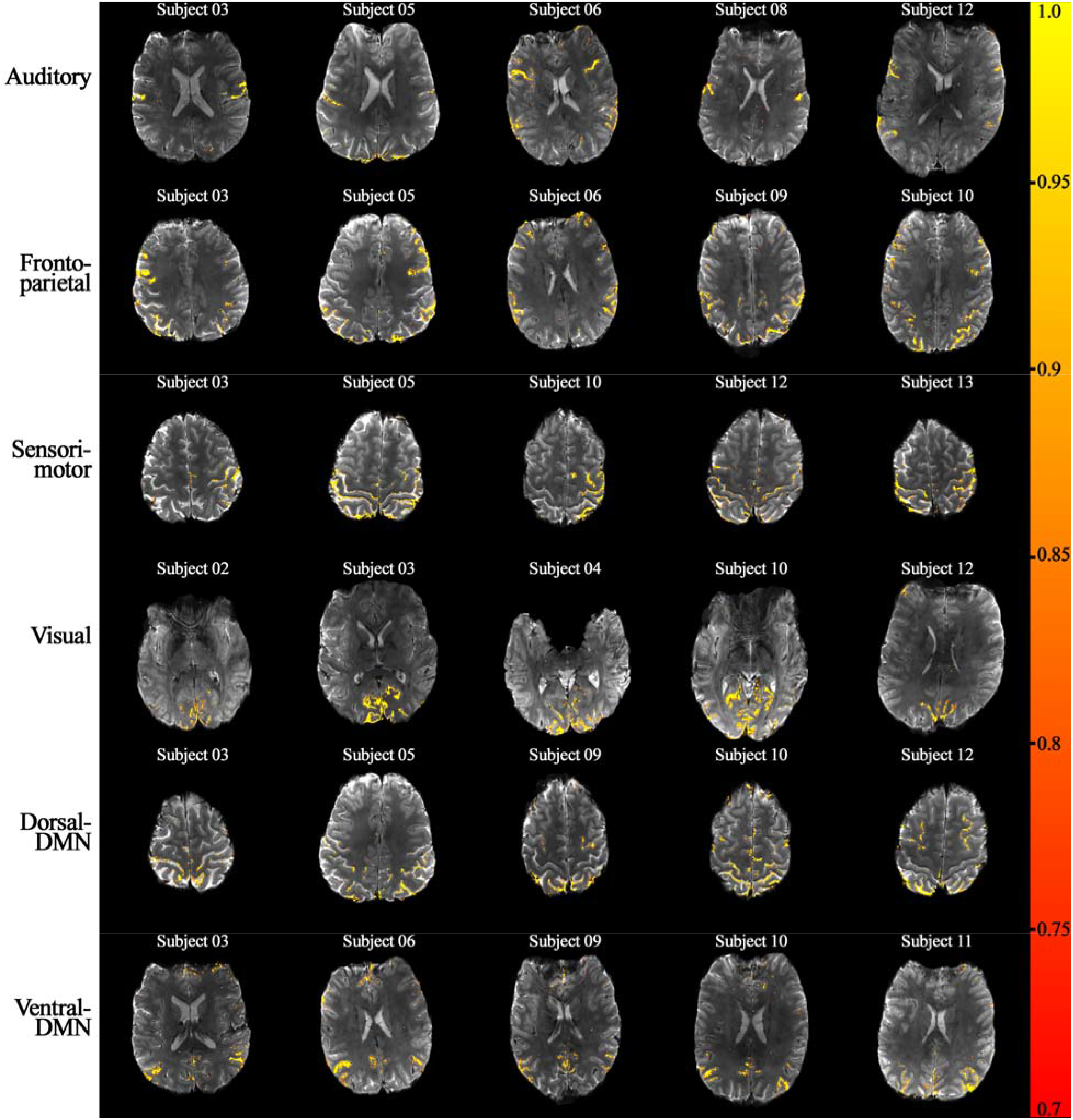
Reproducibility in the detection of resting-state networks. For each of the six resting-state networks (auditory, fronto-parietal, sensorimotor, visual, dorsal-DMN and ventral-DMN), the activated voxels from five representative subjects are shown. A similar activation pattern can be observed over all subjects for each network, demonstrating the reproducibility of functional mapping.

The quality of functional scans from the 13 subjects was assessed by inspecting the image SNR. The averaged image SNR from all subjects for all temporal scans was 251.63, where its standard deviation across subjects was 26.64. Notwithstanding the fact that the SNR varies between subjects, the relatively small standard deviation across subjects indicates that robust fMRI acquisition was performed throughout. Moreover, the standard deviation along the temporal dimension was calculated for each fMRI session, and the averaged value from all subjects was 4.29. This result demonstrates that no significant SNR variation along the temporal dimension was accrued during the fMRI session, which would have otherwise had a deteriorative effect on the quality of the functional data. The SNR results for each subject can be found in S2 Table.

The application of TR-external EPIK for resting-state fMRI was also demonstrated with an isovoxel protocol (0.63 × 0.63 × 0.63 mm^3^) for the same group of subjects as employed in the half-millimetre protocol. These results also show that the activated voxels are well localised along the cortical ribbons, and similar resting-state networks to those observed from the half-millimetre protocol were identified for all subjects. Furthermore, the effect of the correction method for the phase-related venous contribution was also demonstrated using the same protocol. A more detailed description of these results is given in Supporting Information (S3 – S5 Fig).

### 2.4. Imaging performance of TR-external EPIK

In order to provide insight into the spatial resolution and brain coverage provided by TR-external EPIK, its imaging performance was evaluated by directly comparing the imaging parameters employed in several previous submillimetre-resolution fMRI studies [15,17,18,21–25,43,44]. The parameters compared were: (1) the cubic volume of the imaging FOV containing brain tissue; (2) volume of a single voxel; and (3) the number of voxels per temporal volume determined by the imaging matrix size, whereby the first two parameters essentially reflect brain coverage and nominal spatial resolution employed in each study. As shown in Fig 6a, compared to prior publications, TR-external EPIK has the largest volume coverage containing brain tissue (4762.80 cm^3^). Although it can be seen that one other study (Sharoh^2019^) has a very comparable brain volume coverage (4659.48 cm^3^), its single voxel volume (0.80 mm^3^), indicating a much coarser spatial resolution, is much larger than that of TR-external EPIK (0.26 mm^3^)(Fig 6b). As a consequence, the number of voxels per each temporal volume in the study by Sharoh^2019^ et al. (5.82 M) is significantly smaller than that of TR-external EPIK (17.98 M)(Fig 6c).

**Fig 6.**
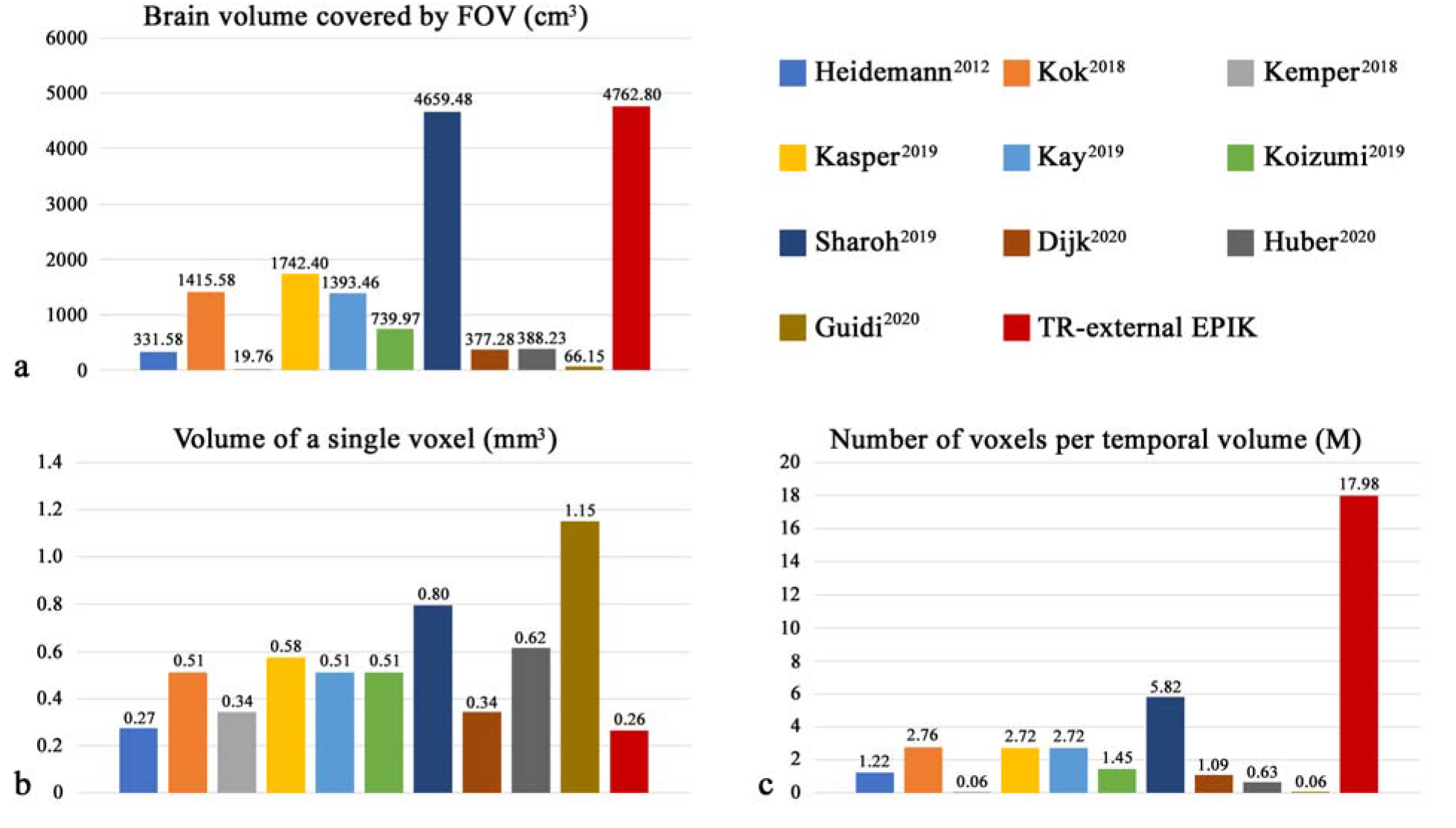
Imaging performance of TR-external EPIK in comparison to other submillimetre-resolution studies. (**a**) Cubic volume of the imaging FOV, (**b**) volume of a single voxel and (**c**) number of voxels per temporal volume. TR-external EPIK shows the largest brain volume (4762.80 cm^3^) and the smallest voxel volume (0.26 mm^3^), which leads to a significantly larger number of voxels per temporal volume (17.98 M).

Although Fig 6b also demonstrates that several other studies have performed their fMRI experiments with a relatively small voxel volume, i.e. approx. 0.6 mm^3^ (cubic root: 0.84 mm), and three studies in particular (Heidemann^2010^, Kemper^2018^ and Dijk^2020^) have shown comparable voxel volumes (0.27, 0.34 and 0.34 mm^3^) to TR-external EPIK (0.26 mm^3^), their corresponding brain coverage (331.58, 19.76 and 377.27 cm^3^) was significantly smaller, by an order of magnitude or more, than that obtained using TR-external EPIK (4659.48 cm^3^).

In other words, the three studies alluded to above were only able to interrogate brain function with high spatial resolution over a very circumscribed part of the brain. Overall, the results in Fig 6 show that TR-external EPIK provides the largest brain volume, while simultaneously providing the highest nominal spatial resolution, which leads to the largest number of voxels per temporal volume. More detailed imaging parameters of each study, such as FOV or in-plane resolution, can be found in Supporting Information where the technical performance of TR-external EPIK is also further assessed in comparison to that of the community standard method, EPI (S3 Table, S4 Table and S6 Fig).

## 3. Discussion

### 3.1. TR-external EPIK and its imaging performance

This work demonstrates the ability of the TR-external EPIK mapping technique to achieve 0.26 mm^3^ resolution and the acquisition of resting-state functional signals with whole-brain coverage at 7T. To achieve a voxel size with a half-millimetre in-plane resolution, a novel fMRI method was developed based on EPIK and a TR-external EPI phase correction.

The method uses high-resolution EPIK to simultaneously achieve both the large number of echoes required and the reduced TE by applying segmented, interleaved sampling for peripheral k-space. The central k-space (keyhole) is fully sampled with Nyquist criteria, which ensures optimum SNR and CNR for each temporal frame, as most information in natural images is distributed around lower frequencies in k-space. The feasibility of using EPIK for fMRI has been verified in several previous publications [32,34,35], where the performance of EPIK was rigorously evaluated in comparison to that of the community standard method, EPI. These prior results clearly demonstrate that the performance of EPIK is superior to that of EPI in terms of spatial resolution and robustness against susceptibility differences while maintaining a comparable temporal SNR and the detection of BOLD signals. The advantages of EPIK for fMRI, in comparison to EPI, have been thoroughly investigated in the earlier works and are not discussed in detail herein.

The TR-external phase correction scheme shortens the required TE by acquiring the navigator echoes in a separated, low-flip-angle excitation loop, which leads to higher resolution fMRI^27^compared to the standard phase correction scheme [45]. In this work, this benefit, in combination with EPIK, allowed an in-plane matrix size of 408 × 408 (0.51 × 0.51 mm^2^) with a TE of 22 ms and a FOV of 210 × 210 mm^2^. For the same TE, the maximum possible matrix size for EPI was determined to be 282 × 282 (0.74 × 0.74 mm^2^)(S3 Table). Importantly, the TE is that typically used for fMRI at 7T for optimal BOLD sensitivity and the FOV was set large enough to cover the entire brain sliced in the axial direction.

There have been numerous high-resolution fMRI studies using a submillimetre voxel size. However, due to the technical limitations of current fMRI techniques, the measurement of resting-state functional signals has not previously been performed with the spatial resolution and brain coverage demonstrate by TR-external EPIK. For the literature comparison shown here, TR-external EPIK outperforms the other methods in terms of the nominal spatial resolution (0.51 × 0.51 × 1.0 mm^3^) and the brain volume (4762.80 cm^3^) covered by the imaging FOV. For most previous methods, submillimetre - resolution was achieved using a rather small FOV (< 200 × 200 mm^2^) and a considerably smaller matrix size (< 200 × 200) than the half-millimetre TR-external EPIK protocol (S4 Table), meaning that only a part of the brain was imaged. Accordingly, TR-external EPIK leads to a much larger number of voxels per temporal volume (17.98 M) when compared to the previous methods, while also having the smallest voxel volume (0.26 mm^3^). Depending on the method selected from the literature, it may be possible to achieve an increased spatial resolution or larger slice coverage by adjusting the imaging protocols. Despite the difficulty of performing a fair comparison between different methods under the given imaging conditions, the results show the clear improvements afforded by TR-external EPIK compared to the previously reported methods.

### 3.2. Spatial resolution of TR-external EPIK

The spatial resolution of TR-external EPIK enabled the identification of mesoscale anatomical structures such as small cerebral vessels or a dark stripe which is located at approximately 50% of the cortical depth in discrete regions throughout the brain. Within the calcarine sulcus, this stripe resembles the stria of Gennari, which is a heavily myelinated tangential band at 60% of the cortical depth [46] and is only visible in high-resolution structural MRI [47–49]. Although the stria of Gennari is a feature unique to the primary visual cortex [50], we also detected comparable stripes in the anterior wall of the postcentral gyrus (primary somatosensory area 3), on the Heschl gyrus (primary auditory cortex), and in the parietal operculum (secondary somatosensory cortex). Furthermore, of these areas, only the primary somatosensory cortex is characterised by the presence of two clearly identifiable Baillarger stripes (myeloarchitectonic layers 4 or 5b), of which, the inner one is more densely myelinated than the outer one [50]. A common feature of all areas containing a dark stripe is the presence of a prominent and densely packed inner granular layer. Therefore, we have interpreted the stripe as being cytoarchitectonic layer IV, and not myeloarchitectonic layer 4 or 5b. It is important to note that these features were visible in fMRI images, which are primarily intended to depict function and not to demonstrate anatomical features.

In EPI-based sequences, the relatively long readout leads to a deteriorated spatial resolution in the reconstructed image, resulting in image blurring. This artefact becomes worse for higher spatial resolution, which usually demands a correspondingly longer readout. The readout length for the half-millimetre TR-external EPIK protocol in this work is 62.01 ms, whereas the length for a comparable EPI protocol with the same matrix size is 135.15 ms. These echo train lengths result in a full-width at half-maximum (FWHM) of 0.0107 and 0.0174 for TR-external EPIK and EPI for the point spread function (PSF) simulation (S6 Fig). Although inspection of an actual EPI scan in comparison to TR-external EPIK was not performed here, the numerical values from the simulation indicate that TR-external EPIK has a better spatial resolution than the EPI method. The higher spatial resolution of EPIK has already been demonstrated for identical imaging parameters to EPI in previous work [32]. In addition to the higher spatial resolution, the use of relatively small acceleration factors for parallel imaging and multi-band techniques in this work (i.e. three-fold each) ensured reliable image reconstruction without inducing significant intra-slice aliasing or inter-slice leakage artefacts [40], which can sometimes be caused by these acceleration techniques. Further improvements in spatiotemporal resolution may also be possible in TR-external EPIK by means of a higher acceleration factor (? 4), as demonstrated in previous works [51,52].

### 3.3. Resting-state networks

For application purposes, the half-millimetre protocol was verified with resting-state fMRI on eighteen healthy subjects. A number of resting-state networks distributed throughout the brain were reliably identified from a single fMRI session for all subjects except one, who exhibited relatively large head displacements during the fMRI scans. From the data set of a representative subject, localised mapping along the cortical ribbon is shown with the six selected resting-state networks (dorsal-DMN, ventral-DMN, visual, sensorimotor (LH), sensorimotor (RH) and fronto-parietal). Here, a line profile of the BOLD signals additionally showed signal variation within the cortical regions, indicating that TR-external EPIK may be useful for cortical depth-dependent functional studies. The activated voxels were displayed and directly overlaid on the mean image of the re-aligned functional scans rather than using an anatomical scan, which is typically acquired with MPRAGE. In this study, co-registration of functional scans to the anatomical scan or spatial normalisation to a common template was not performed. This was to prevent any possible deleterious effects, which might affect the localisation of functional signals, from being introduced by these steps, i.e., all analyses were performed in subject space. Interestingly, the very high spatial resolution of the functional scan demonstrated in this work (Fig 1) suggests that the TR-external EPIK image could be directly used for anatomical reference. A recent work has presented a method that acquires the T_1_-contrast anatomical scan with the same base sequence used for the functional scan [53]. This approach effectively reduces co-registration errors stemming from the use of different imaging sequences for functional (e.g. EPI) and anatomical imaging (e.g. MPRAGE). The same approach can also be applied to our method, and hence, it is expected that TR-external EPIK with T_1_-contrast could be used as a substitute for MPRAGE. Implementation of the technique and its application in fMRI will be demonstrated in future studies.

The detection of resting-state networks from 13 subjects verified the robust use of TR-external EPIK for resting-state fMRI. The reproducibility analysis further showed that each of six well-known resting-state networks (auditory, fronto-parietal, sensorimotor, visual, dorsal-DMN and ventral-DMN) was characterised with a similar activation pattern across the subjects. The SNR results also demonstrated that the functional images were acquired without any significant SNR variation during each fMRI session, and inter-subject SNR variation was also shown to be relatively small.

In this work, typical resting-state fMRI acquisitions were performed with an isovoxel size (0.63 × 0.63 × 0.63 mm^3^). The resting-state fMRI results from this protocol also showed that resting-state networks can be reliably detected in the same 13 subjects with a high mapping fidelity onto the cortical ribbon, verifying the use of TR-external EPIK for isotropic voxel applications.

### 3.4. Non-BOLD contrasts

In order to overcome the relatively low spatial specificity to neuronal activation in gradient-echo (GE) BOLD, several non-BOLD contrasts have previously been demonstrated. Non-BOLD contrasts probe the functional locations more directly than the BOLD-contrast, for instance, using the neuronal signal changes generated by cerebral blood flow, cerebral blood volume or cerebral metabolic rate of oxygen [54]. Exemplary methods exploiting each of the aforementioned contrasts are ASL [55], VASO [8] and calibrated BOLD [56], respectively. It has been shown that these contrasts can offer improved spatial specificity and functional quantification compared to GE-BOLD contrast, and hence, can deliver more direct physiological responses for cortical depth-dependant activations [54]. However, due to the relatively low sensitivity of these contrasts (10%-20% for ASL, 40%-60% for VASO and 5%-15% for calibrated BOLD when compared to GE-BOLD) [54], their use for submillimetre-resolution fMRI is still challenging. Furthermore, the underlying mechanisms of each contrast (e.g. labelling, blood nulling or reach of functional steady-state) usually requires a much longer TR than the GE-BOLD method, which may significantly hinder whole-brain fMRI for a given TR. For these reasons, fMRI methods with GE-BOLD contrasts are still widely used in submillimetre-resolution fMRI. Previously, there have been several approaches to reduce the large-vessel BOLD effect in GE-BOLD by applying a phase correction method [41,42] or replacing it with spin-echo BOLD. Both approaches can, in principle, be applied to TR-external EPIK. In this work, the phase correction method [41,42] was applied to the functional data sets to partially minimise the influence of large cerebral vessels. In an earlier work, a spin-echo version of EPIK was also verified for the detection of haemodynamic signals in perfusion studies [37]. This suggests that a spin-echo configuration can be systematically transferred to TR-external, gradient-echo EPIK and the same high-resolution advantage obtained in this work can also be expected in its spin-echo version, resulting in increased spatial specificity to aid the detection of neural activity.

### 3.5. Functional resolution

One of the main challenges of interpreting high-resolution fMRI relates to the complexity of the neural hemodynamic responses and the heterogeneous vascular network topology of the cerebral cortex [57,58]. Neuronal activity is coupled to an increase in the cerebral metabolic rate of oxygen, which, under conditions of normal neuro-vascular coupling, triggers a local increase in the cerebral blood flow and volume that can be detected with fMRI sequences. However, the specific temporal dynamics of the vascular responses (i.e., the hemodynamic response function) can alter depending on the vascular environment, e.g. across different cortical layers [59,60]. In particular, the heterogeneously distributed cerebral veins and venules (large pial veins running tangentially to the cortex and ascending venules draining blood from deep cortical layers towards the surface) introduce a bias in the fMRI signal, especially in GE-studies where the contribution of the macro-vasculature is considerably higher. Moreover, the varying capillary density across the cortical depth further conditions the actual spatial resolution of functional responses. Simulation studies have demonstrated an enhanced specificity of the fMRI signal towards the micro-vasculature at higher fields using short TEs [61]. Additionally, several works have estimated the PSF of the hemodynamic responses linked to neuronal activity, which would represent the minimum space unit that could be resolved with fMRI contrast. This ultimate resolution was estimated to be 0.1 mm in an optical study of the cerebral blood volume in mice [62], 0.86 mm for spinecho BOLD fMRI, and 0.99 mm for GE-BOLD fMRI in a human study of ocular dominance columns [63]. Taking into account the unavoidable presence of physiological motion and the resultant blurring in human fMRI, the true haemodynamic PSF is probably narrower than the PSF in the presence of physiological motion, which in turn results in a higher putative functional resolution. Notwithstanding the controversial definition of the ultimate functional spatial resolution, the fact that published, experimental submillimetre fMRI studies have detected distinct signals originating in highly segregated cortical units, such as cytoarchitectonically-defined layers (e.g., cortical depth-specific activation upon engagement of a finger in a sensory or a motor task [8]), strongly supports the ambition of acquiring and analysing functional data with submillimetre-resolution as high as provided here by TR-external EPIK (0.51 × 0.51 × 1.0 mm^3^; 0.63 × 0.63 × 0.63 mm^3^).

### 3.6. Potential use of TR-external EPIK

In the fMRI community, whole-brain coverage is very often demanded as one of the first requirements for fMRI. TR-external EPIK is therefore expected to be deployable in many types of functional studies requiring whole-brain as well as high-resolution mapping. In addition, although it was not demonstrated here, EPIK is able to alleviate the geometric distortions caused by the susceptibility differences around the tissue boundaries, when compared to EPI [30,32,34,35]. Hence, TR-external EPIK is also expected to be beneficial for exploring brain regions such as the frontal or temporal lobes, in which susceptibility artefacts are typically seen with standard EPI methods. Moreover, the substantial reduction in the minimum TE in TR-external EPIK enables its use in a very straightforward way at a higher field strength (e.g. 9.4T), where T_2_* decay is faster and therefore the BOLD-optimised TE is shorter. In event-related task fMRI, the haemodynamic response changes are, in general, not as smooth as those from a resting-state or block-based paradigm [34]. Consequently, the functional response may require EPIK to update the peripheral k-space more quickly. In light of this, the use of EPIK for event-related fMRI needs to be carefully investigated in future work.

## 4. Materials and Methods

### 4.1. TR-external EPIK

Fig 7a shows a sequence diagram of the proposed TR-external EPIK method. The entire TR loop consists of two excitation loops. In the first excitation loop (α_PC_; TR_PC_), three navigator echoes are acquired with a relatively small flip angle to avoid strong signal saturation in the next excitation loop. However, in order to yield a sufficient SNR for the navigator echoes, the angle should not be too small. In this work, 9° was employed for α_PC_, as demonstrated by previously [27]. The second excitation loop (α_Main_; TR_Main_) presents a typical echo-planar acquisition whereby the data sampling is performed according to the EPIK scheme (Fig 7b). It should be noted that the lack of navigator echoes in the second loop facilitates a decreased minimum TE when compared to a counterpart EPI sequence. This benefit, in turn, makes it possible to further increase the imaging matrix size for any given TE for fMRI purposes.

**Fig 1.**
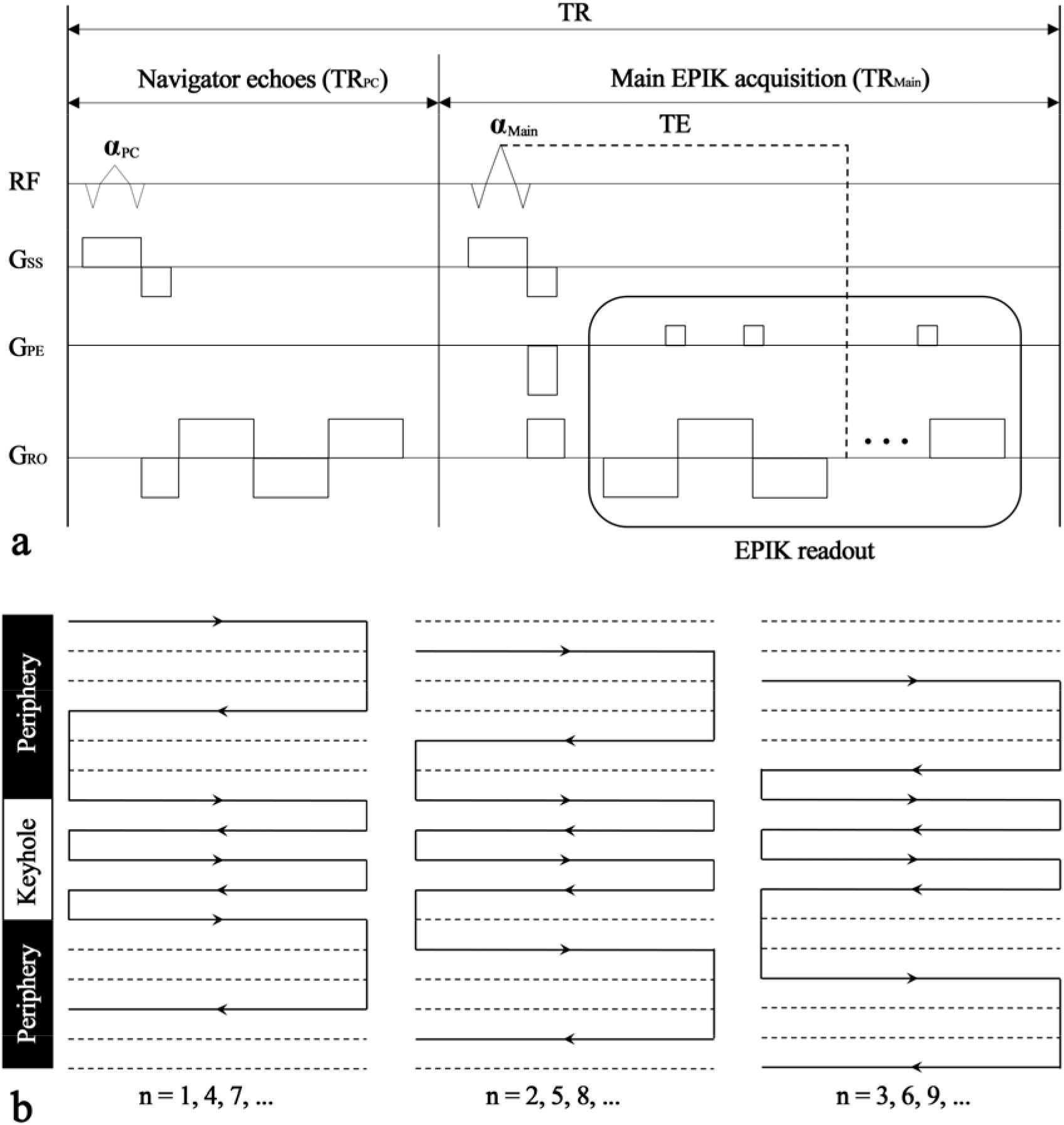
EPIK with TR-external EPI phase correction (TR-external EPIK). (**a**) Schematic representation of TR-external EPIK and (**b**) detailed description of k-space sampling in EPIK. Each TR of the TR-external EPIK consists of a low-flip angle excitation loop (α_PC_; TR_PC_) for navigator echoes and a main excitation loop (α_Main_; TR_Main_) for EPIK readout (outlined by the rectangle in a). The EPIK scan is acquired according to the k-space trajectories shown in b, in which the solid and dashed lines indicate lines to be sampled and to be skipped, respectively. The first k-space line acquired in the 1st, 2nd and 3rd scans (i.e. n = 1, 2 and 3) is 1, 2 and 3, respectively. The first line acquired becomes 1 again for the 4th scan (n = 4), in which the original line 1 is discarded ensuring a refreshed periphery and thereby avoiding autocorrelations in the time-series data used for fMRI analyses.

Fig 7b illustrates a schematic representation of three-shot EPIK, with respect to the temporal indices. Each measurement consists of full Nyquist-rate sampling (Δk_y_ = 1/FOV) of the central k-space region and sparse sampling (Δk_y_’ = 3/FOV) of the peripheral k-space regions. As shown in the figure, sparse sampling is performed as in multi-shot EPI and the missing k-space lines for every time frame are reconstructed with a sliding-window reconstruction scheme [64]. In this way, EPIK can achieve a higher *apparent* temporal resolution than single-shot EPI. In contrast to multi-shot EPI, the complete sampling of central k-space, referred to here as the “keyhole”, ensures an optimum SNR and CNR for every single time frame. In this work, 48 central k-space encoding lines were configured as the keyhole region [38,39]. It is important to note that the peripheral k-space used in each image is updated every three shots to prevent the introduction of autocorrelations into the time-series data used for the fMRI analyses.

The half-millimetre protocol in this work was implemented based on the above TR-external EPIK scheme in combination with other typical acceleration techniques, such as partial Fourier [65], parallel imaging [66] and multi-band techniques [67]. The detailed imaging parameters were as follows: TR = 3500 (i.e. 15.7 ms for TR_PC_ + 3484.3 ms for TR_Main_), TE = 22 ms, FOV = 210 × 210 mm^2^, matrix = 408 × 408 × 105 slices (0.51 × 0.51 × 1.0 mm^3^), partial Fourier = 5/8, 3-fold in-plane/3-fold inter-plane (multiband) acceleration, bandwidth = 721 Hz/Px and αPC/αMain = 9°/90°. In addition, an isovoxel protocol was also implemented with a matrix size of 336 × 336 × 123 slices (0.63 × 0.63 × 0.63 mm^3^) and bandwidth = 875 Hz/Px, while other conditions were kept identical. The above imaging configuration was employed on a Siemens Magnetom Terra 7T scanner with a single-channel Tx/32-channel Rx Nova medical coil supplied by the manufacturer.

### 4.2. In vivo measurements

All *in vivo* measurements in this work were performed on healthy volunteers screened, in addition, for neurological or psychiatric illnesses. After a complete description of the study, written informed consent was obtained before scanning. The local institutional review board (RWTH Aachen University, Germany) approved the study protocol (EK 346/17), screening questionnaires, and consent forms. Eighteen healthy volunteers (14 males, 4 females; mean age, 30.22 years; range, 23-47 years) were recruited in the study. In order to control for the effect of physiological noise, respiratory and cardiac signals were recorded using a pneumatic belt positioned around the subject’s chest and a pulse-oximeter placed on the 2nd, 3rd or 4th finger of the left hand.

### 4.3. Resting-state fMRI and data analysis

Resting-state fMRI was performed using both the half-millimetre and the isovoxel protocols described above. The data acquisition consisted of four dummy scans, in order to reach a steady-state, and 172 scans to acquire 10-minutes of resting-state fMRI data. Functional data were pre-processed using AFNI (Analysis of Functional NeuroImages, NIH, Bethesda, MD) to perform slice-timing correction, realignment, regression of cardio-respiratory signals, regression of the mean white-matter and mean CSF signal, temporal filtering with regression of motion parameters, and correction of the phase-related venous effects [41,42]. Data sets from five subjects were excluded due to the following reasons: one subject’s data set showed extremely low functional activation of the reported networks (possibly linked to relatively high motion displacement from the subject), two subjects had very noisy signals in the recorded physiological data (i.e. respiratory and cardiac signals) leading to unreliable data pre-processing, and the other two subjects had systematic difficulties causing the physiological data to only be partially recorded. Independent component analysis was performed on the functional data with Melodic, in FSL (FMRIB Software Library, Oxford, UK), specifying a number of components equal to 80, which resulted in 80 maps where the probability at each voxel quantified the probability of the voxel of belonging to a given network (i.e., component characterised by a certain time course). From the 80 components, the best matching component to present each resting-state network (i.e. dorsal-DMN, ventral-DMN, visual, sensorimotor (LH), sensorimotor (RH) and fronto-parietal) was manually selected and the activated voxels were obtained with a statistical threshold (probability ≥ 0.7). Here, for illustrative purpose, clusters containing less than three voxels were excluded from the visualisation. The line profiles of BOLD and background image signals in Fig 3 were computed using an internal function (‘improfile’) of MATLAB (MathWorks Inc.). The lines to be examined were selected such that they cross the cortical ribbon perpendicularly. The length of the examined line was 10 voxels (5.1 mm), and 100 points were sampled along its length. The GM regions were obtained by applying the standard segmentation routine in FSL (FAST; T_2_-weighted) to the mean image of the re-aligned functional scans. The analysis of the line profile was performed for the three resting-state networks: dorsal-DMN, sensorimotor for the right hemisphere and visual.

### 4.4. 3D visualisation

The 3D rendered surface of GM and WM (Fig 1) was obtained using a 3D visualisation tool, 3D Slicer 4.10.2 (www.slicer.org). The activated voxels from the six resting-state networks (i.e. dorsal-DMN, ventral-DMN, visual, sensorimotor (LH), sensorimotor (RH) and fronto-parietal) were projected onto the surface and displayed with 50% transparency for visualisation purposes (Fig 2).

## Author Contributions

N.J.S. proposed EPIK for laminar fMRI research; S.Y. designed the TR-external EPIK protocol and programmed the sequence; S.Y. developed the corresponding image reconstruction software; S.Y. performed phantom experiments; S.Y. and P.P. performed in vivo experiments. S.Y. performed phantom and *in vivo* data analysis; S.Y. wrote the first draft of the manuscript including all the figures and tables and critically revised it; P.P. performed resting-state fMRI data analysis and wrote the corresponding texts; N. P.-G. delineated anatomical structures and wrote the corresponding texts. All authors reviewed the manuscript. N.J.S. invented and patented the original EPIK sequence; N.J.S. critically reviewed the final version of the manuscript and takes responsibility for its authenticity.

