## Supplementary Information for "Mapping of Whole-Brain Resting-State Networks with Half-Millimetre Resolution"

**Supporting Information**

**Verification of TR-external EPIK for submillimetre voxel sizes**

TR-external EPIK was verified with various submillimetre voxel sizes. The imaging parameters of each protocol are shown in S1 Table. With a fixed FOV (210 × 210 mm^2^) and TR/TE (3500/22 ms), several isovoxel protocols were configured to yield a nominal spatial resolution of 0.88, 0.73, 0.63, 0.58 and 0.55 mm (i.e. voxel volumes of 0.68, 0.39, 0.25, 0.20, 0.17 mm^3^; 2nd to 6th columns). For each case, the matrix size employed and the in-plane acceleration setting are also shown in the table. The last row of the table shows the overall in-plane acceleration factor achieved by each protocol, which is computed by the target matrix size divided by the number of phase encoding lines encoded. As shown in the table, the in-plane acceleration factor becomes larger for higher nominal spatial resolution, and, for the 0.55 mm case, the acceleration factor reaches as high as 10.67. The number of keyhole lines in each protocol was determined such that the overall number of phase encoding lines to be encoded should be an integer value for the given parallel imaging and partial Fourier acceleration conditions. A more detailed explanation of the k-space sampling in EPIK is described in the ‘Methods’ section.

For the five isovoxel protocols described above, the quality of the reconstructed image and the image SNR were inspected using data sets obtained from a uniform spherical phantom and a healthy subject. Here, every protocol was additionally accelerated using the same multi-band condition (*R* = 3) as used in the actual fMRI measurements in this work. S1 Fig shows reconstructed phantom and *in vivo* images for a representative slice location. All images can be seen to be well reconstructed without any severe image degradations, clearly demonstrating and validating the use of TR-external EPIK for submillimetre voxel sizes. The reconstructed images do not show any significant N/2 ghost artefacts or aliasing artefacts that can sometimes occur due to EPI phase correction and the unfolding process from parallel imaging or multi-band techniques. In addition, the enhancement of spatial resolution can also be observed as the matrix size increases. The image SNR of each protocol is displayed at the bottom of each image. As can be visually verified, the SNR decreases as the nominal spatial resolution increases.

| **Resolution (mm^2^)** | **0.88 × 0.88** | **0.73 × 0.73** | **0.63 × 0.63** | **0.58 × 0.58** | **0.55 × 0.55** | **0.51 × 0.51*** |
| --- | --- | --- | --- | --- | --- | --- |
| **Matrix** | 240 × 240 | 288 × 288 | 336 × 336 | 360 × 360 | 384 × 384 | 408 × 408 |
| **Keyhole lines** | 60 | 54 | 48 | 54 | 42 | 48 |
| **Parallel Imaging** | 2 | 3 | 3 | 3 | 3 | 3 |
| **Partial Fourier** | 5/8 | 5/8 | 5/8 | 5/8 | 5/8 | 5/8 |
| **Factor of in-plane Acceleration** | 5.33 | 9.00 | 9.88 | 9.73 | 10.67 | 10.46 |

**S1 Table. TR-external EPIK protocols for several submillimetre resolutions.** All the protocols were configured under the same FOV (210 × 210 mm^2^) and TR/TE (3500/22 ms) conditions. The slice thickness of each protocol was adjusted to yield an isovoxel size, except the half-millimetre protocol (marked with an asterisk) in which 1 mm thickness was used for a reasonable SNR. The in-plane acceleration factor, achieved by the half-millimetre protocol, was around 10.46, which is significantly larger than those (4.8) when EPI is used instead of TR-external EPIK.

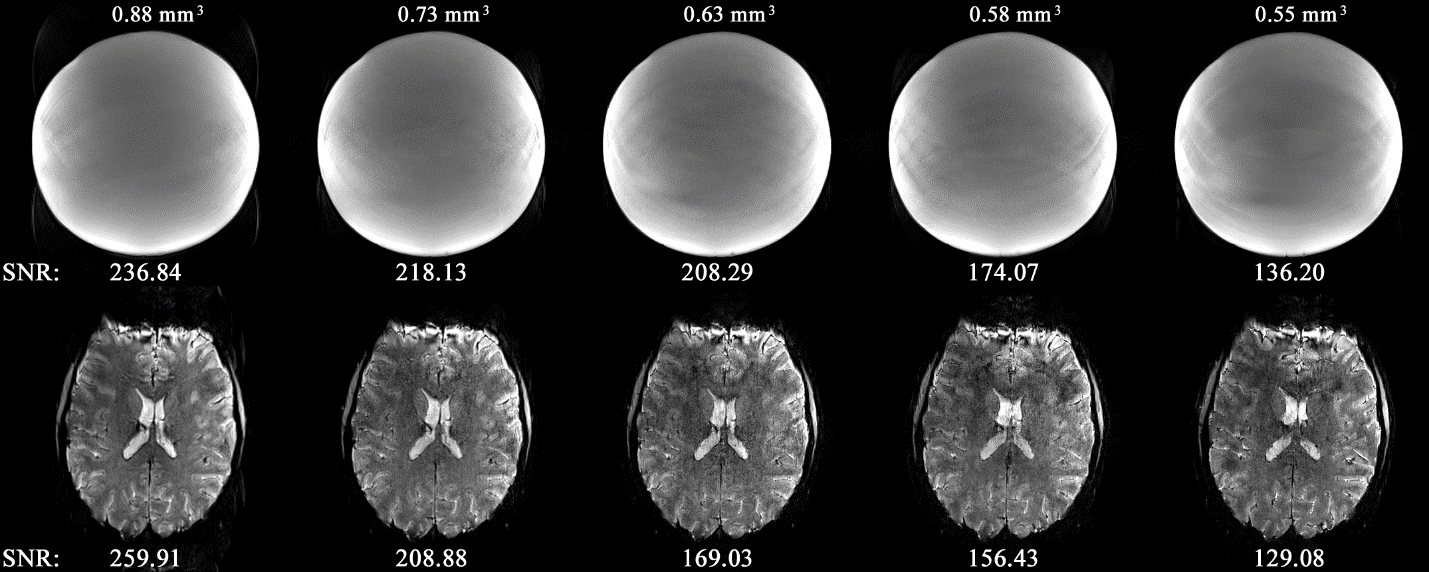

**S1 Fig**. **TR-external EPIK images for several isovoxel submillimetre protocols.** The top and bottom rows show images obtained from a spherical phantom and an *in vivo* data set from a healthy subject. All the images are well reconstructed, without any significant image degradation such as ghost or image reconstruction artefacts. Despite the substantial reduction of SNR with the increase in matrix size, reliable image reconstruction was achieved for all the protocols.

**Configuration of a protocol with a half-millimetre in-plane resolution**

Under the same FOV and TR/TE condition used in the above submillimetre-resolution protocols, an imaging matrix size of 408 × 408 was possible, yielding a nominal spatial resolution of 0.51 × 0.51 mm^2^. The slice thickness of this protocol was set up such that a reasonable SNR is provided for fMRI. Visual inspection of the above submillimetre-resolution *in vivo* images (S1 Fig) suggests that the SNR delivered by the 0.58 mm and the 0.55 mm protocol is relatively low, although the resultant images were successfully reconstructed. Thus, for more robust fMRI, a slice thickness of the half-millimetre protocol was determined so that its voxel volume is similar to that of the 0.63 mm protocol. Accordingly, the slice thickness was adjusted to 1.0 mm, yielding a voxel volume of 0.26 mm^3^. As described in the main text, this protocol achieved a maximum of 108 slices for the given TR (3500 ms). S2 Fig shows the entire set of axial slices obtained from the half-millimetre protocol. It can be seen that all slices were well reconstructed and do not suffer from any significant image degradation, demonstrating the robustness of TR-external EPIK in generating the nominal 0.26 mm^3^ resolution images.

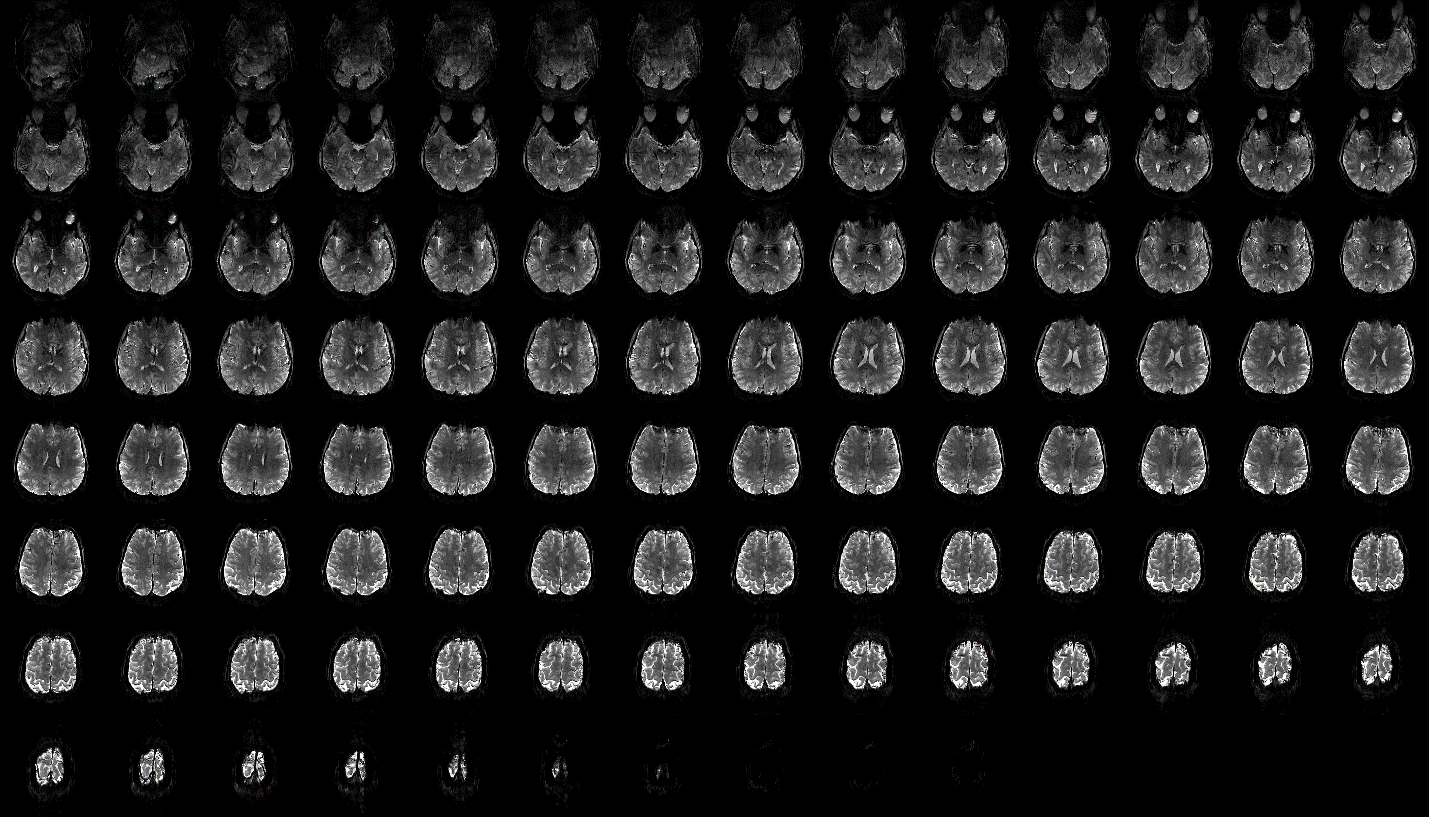

**S2 Fig.** Reconstructed images of the half-millimetre protocol (voxel volume = 0.26 mm^3^) for the entire 108 axial slices.

**Signal-to-Noise Ratio of the half-millimetre protocol**

Image SNR was calculated for the 13 subjects whose resting-state network results are shown in Fig. 4. The calculation was performed for each temporal frame of each subject (i.e. 172 temporal frames per subject). The mean ± SD SNR across the temporal dimension was computed, and the value obtained from each subject is shown in S2 Table. It can be seen that the standard deviation of SNR across the temporal dimension is very small (i.e. in average, 4.3), demonstrating that SNR variation during the fMRI acquisition was not significant for any of the subjects. The averaged SNR values for all subjects and for all temporal volumes were computed as 251.63. The averaged standard deviation of SNR across the 13 subjects was 26.64, indicating that the inter-subject SNR variation was also relatively small.

| **Subject** | **Image SNR (mean ± SD)** |
| --- | --- |
| 1 | 272.88 ± 4.35 |
| 2 | 233.83 ± 4.51 |
| 3 | 289.65 ± 6.62 |
| 4 | 244.25 ± 3.99 |
| 5 | 287.95 ± 4.81 |
| 6 | 279.82 ± 3.24 |
| 7 | 261.40 ± 2.89 |
| 8 | 256.55 ± 2.71 |
| 9 | 256.70 ± 9.07 |
| 10 | 218.67 ± 3.81 |
| 11 | 239.45 ± 3.30 |
| 12 | 220.99 ± 2.87 |
| 13 | 209.00 ± 3.65 |
| **Average** | **251.63 ± 4.29** |

**S2 Table. Image SNR of the half-millimetre protocol.** For each subject, mean ± SD across the temporal dimension (i.e. in total, 172 temporal volumes) is shown. The last row shows the average of the computed values from all subjects.

**Resting-state fMRI with an isovoxel protocol**

The application of TR-external EPIK for resting-state fMRI was demonstrated using the isovoxel protocol (0.63 × 0.63 × 0.63 mm^3^), which had already been verified to yield artefact-free reconstructed images and a reasonable SNR, as shown in S1 Fig. S3 Figa shows the results for the same six resting-state networks examined in the half-millimetre protocol. Similar to that, the activated voxels are well localised along the cortical ribbons. S3 Figb depicts the activated voxels from the six resting-state networks presented on the same background images, demonstrating brain coverage as well as the mapping fidelity onto the grey matter.

S4 Fig is a grid plot summarising the identified resting-state networks obtained from the 17 subjects. The data set from one subject was excluded due to a high mean head displacement; the same subject was also excluded in the half-millimetre protocol case for the same reason. The grid plot shows that the six resting-state networks (auditory, fronto-parietal, sensorimotor, visual, dorsal-DMN and ventral-DMN) were also reliably detected for all of the subjects in the isovoxel data sets.

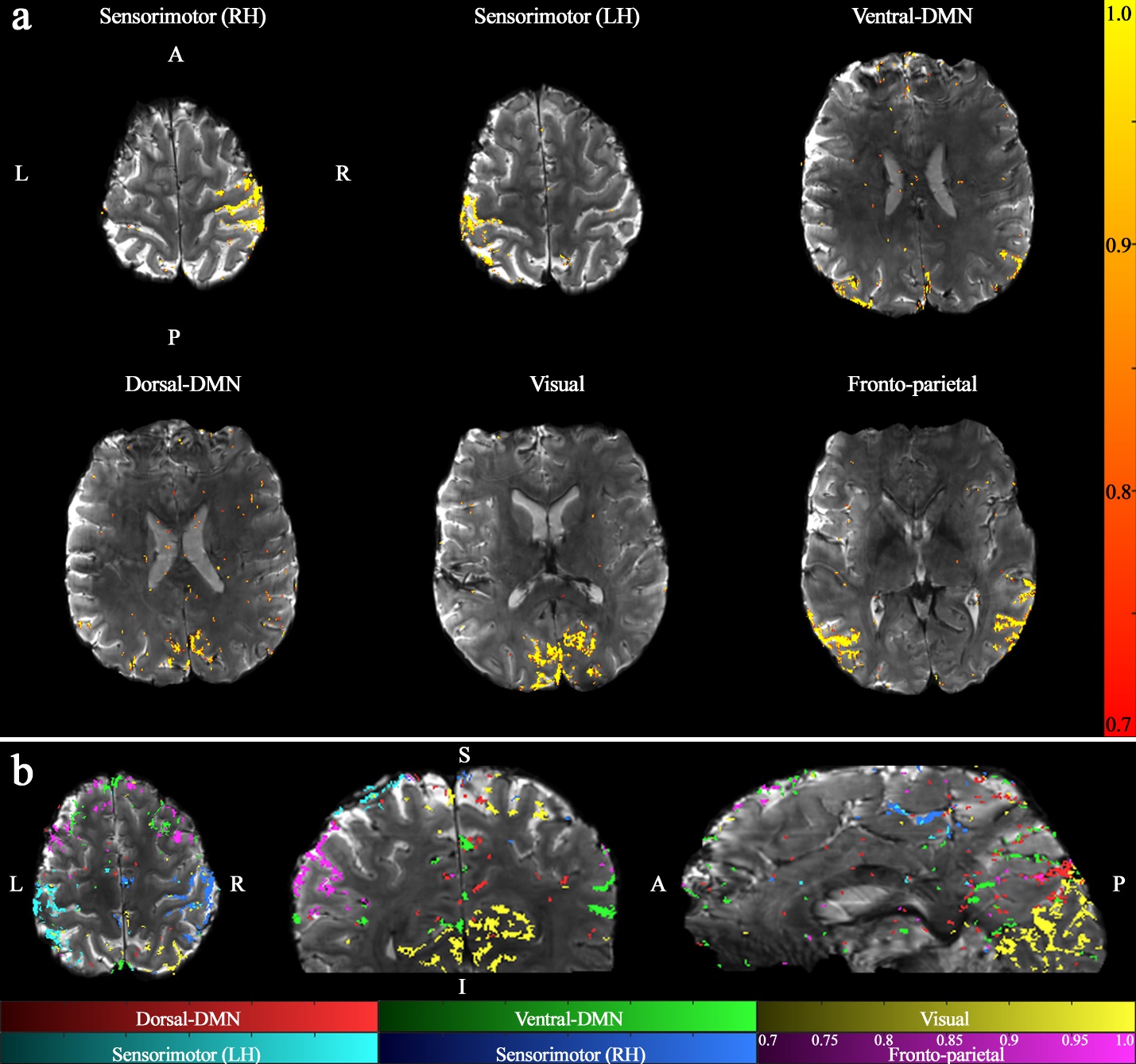

**S3 Fig. Results of resting-state networks obtained with an isovoxel size (0.63 × 0.63 × 0.63 mm^3^).** (**a**) resting-state results for the six selected networks: dorsal-DMN, ventral-DMN, visual, sensorimotor (LH), sensorimotor (RH) and fronto-parietal and (**b**) the six resting-state networks, presented on the same background images in three different sectional views. In panel a, the significance of functional activation is presented with a red-to-yellow scale, and corresponding scale values (i.e. probability) are shown on the right side.

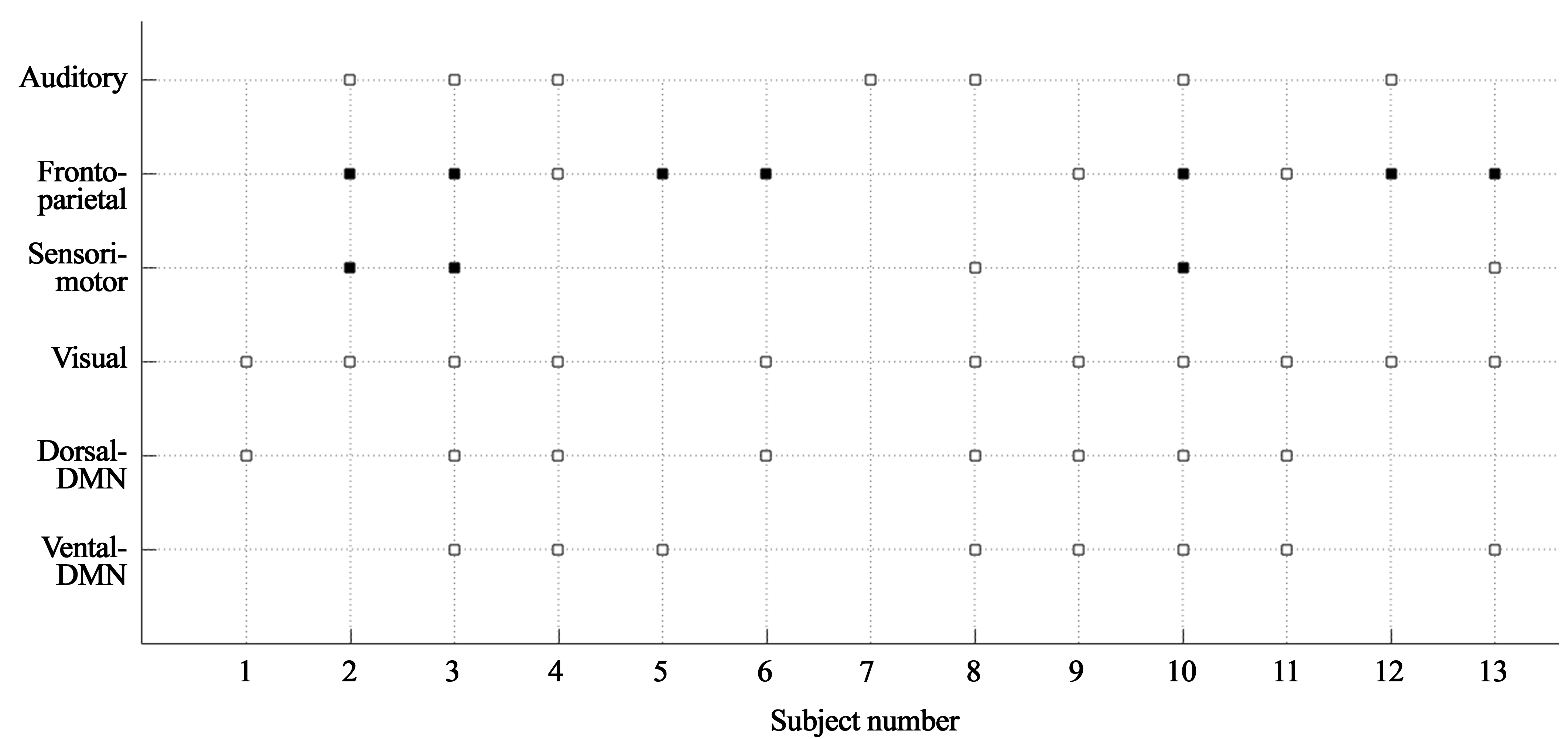

**S4 Fig. A grid plot of resting-state networks with respect to the 13 subjects (0.63 × 0.63 × 0.63 mm^3^ protocol).** For six well-known functional networks (auditory, fronto-parietal, sensorimotor, visual, dorsal-DMN and ventral-DMN), the detected resting-state networks in each subject are marked with a square. The solid squares in the somatosensory and the parietal networks denote the presence of bi-hemispheric activation; an open square is used for the uni-hemispheric activation (either left or right). Out of the 18-subject data sets, a data set that showed relatively high motion was excluded due to very little functional activation. This figure depicts that the resting-state networks were reliably detected for all 17 subjects.

**The effect of pre-processing steps on the functional activation**

As described in the ‘Methods’ section, the resting-state fMRI data underwent several pre-processing steps. Here, in addition to the application of typical procedures such as slice-timing correction, and realignment, the following additional steps were also included: regression of cardio-respiratory signals, temporal filtering with regression of motion parameters and the correction of the phase-related venous effects. In order to inspect the effect of these three steps on the identification of functional signals, the functional maps obtained from each respective stage were compared to those obtained by applying only the first two steps (i.e. slice-timing and realignment). For this comparison, three healthy volunteers who participated in the resting-state fMRI measurements were additionally employed for finger-tapping fMRI. The employed block-paradigm consists of 12 cycles of baseline-activation states, each lasting for six volume acquisitions (6 TRs task – 6 TRs rest). The additional acquisition of finger-tapping fMRI was intended to derive similar functional locations (i.e. motor) for the three subjects, enabling a more straightforward comparison. S5 Fig shows the results obtained with a statistical threshold, p < 0.005. As shown, the reduction of functional activation, especially in non-GM voxels, can be verified at the 2nd stage (regression of cardio-respiratory signals; 2nd row of the figure). This stage accounts for potential false-positive activation created by the signal fluctuations of cardiac and respiratory movements. Temporal filtering and motion regression (3rd row) further diminish the number of activated voxels, as they reduce potential low-frequency noise derived from scanner hardware instabilities and residual motion-related effects in the time course. Finally, in the correction of phase-related venous effects (4th row), a similar reduction of activation can also be observed, particularly around the veins. For the results shown in the figures in the main body of the paper, all five pre-processing steps were used.

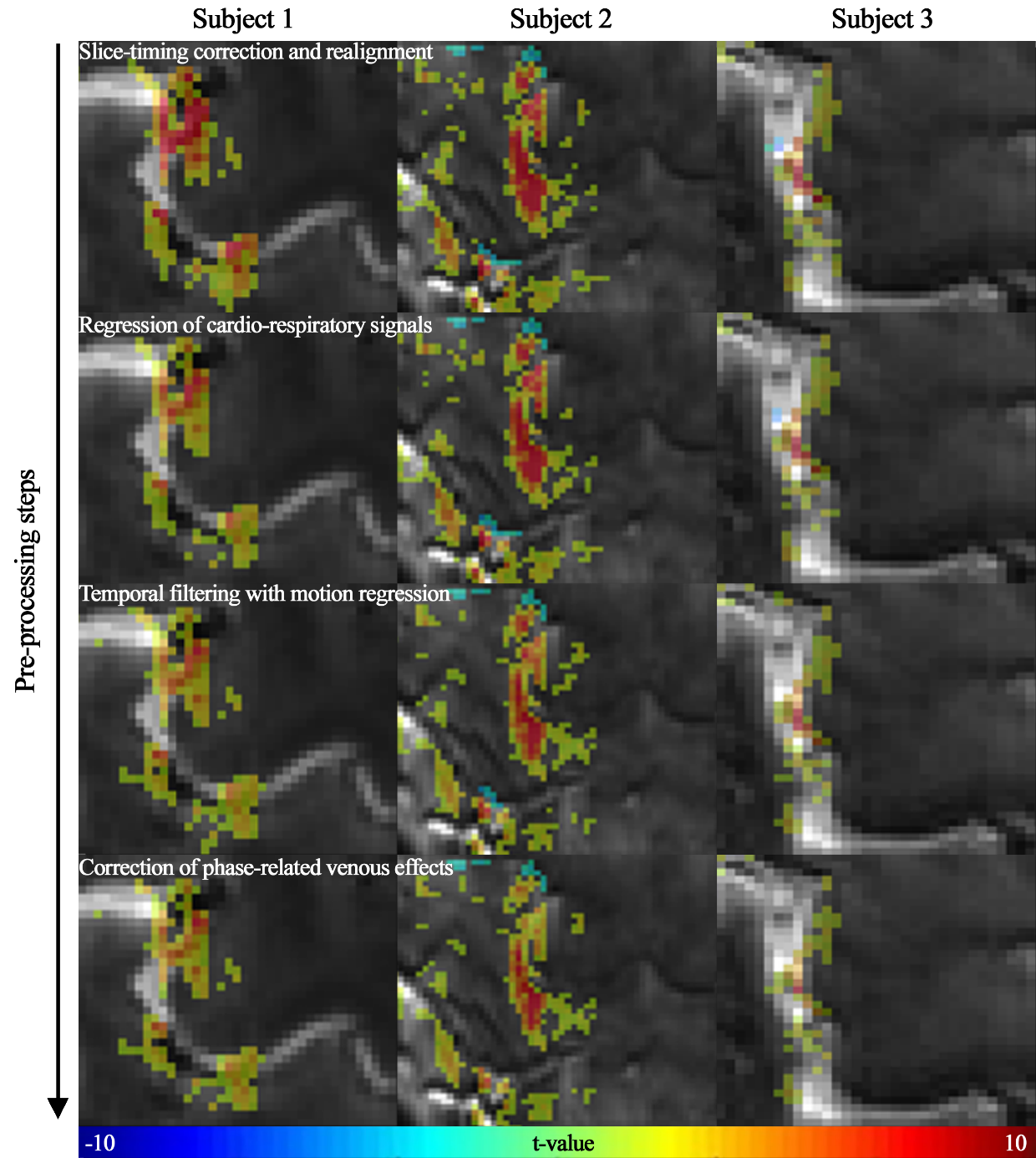

**S5 Fig**. **The effect of the pre-processing steps on the functional signals.** Each row of the figure shows functional voxels obtained at the respective stage of the pre-processing steps: slice-timing correction and realignment, regression of physiological signals (cardio-respiratory motions), temporal filtering with regression of motion parameters and correction of phase-related venous effects. The results are shown for three subjects, acquired with a block-paradigm for finger-tapping using the isovoxel protocol (0.63 × 0.63 × 0.63 mm^3^). Reduced functional activation can be clearly seen in the 2nd and 4th stages (i.e. 2nd and 4th rows).

***Numerical assessment of TR-external EPIK***

The performance of TR-external EPIK was assessed in comparison to that of the community standard method, EPI. The assessment was carried out by inspecting the maximum throughput of each method for the given imaging conditions. As demonstrated in the main text, TR-external EPIK enabled an imaging matrix size of 408 × 408 for a TE of 22 ms. For the same matrix size, the minimum TE required by TR-external EPIK and EPI was 21.53 ms and 36.61 ms, respectively. The minimum TE required by EPI is about 70.04% longer than the TR-external EPIK case, which indicates that optimal BOLD contrast will not be achieved with EPI under these constraints at 7T. When the minimum TE is employed in each method, the minimum TR required to encode a single slice is 158.61 ms and 95.71 ms for EPI and TR-external EPIK, respectively. The minimum TR subsequently results in the acquisition of 66 slices for EPI and 108 slices for TR-external EPIK for a TR of 3500 ms and a multi-band acceleration factor of three (S3 Table). In contrast, when the same TE of 22 ms is mandated, the maximum matrix size achievable with EPI is 282 × 282, yielding a nominal spatial resolution of 0.74 × 0.74 mm^2^. This is a performance degradation of around 31% in both frequency and phase encoding directions when compared to TR-external EPIK (408 × 408; 0.51 × 0.51 mm^2^).

|  | **EPI** | **TR-external EPIK** |
| --- | --- | --- |
| **For the same TE (22 ms)** | **Maximum matrix size achieved** | |
|  | 282 × 282  (0.74 × 0.74 mm^2^) | 408 × 408  (0.51 × 0.51 mm^2^) |
| **For the same matrix size (408 × 408)** | **Min TE/Min TR required** | |
|  | 36.61 ms/ | 21.53 ms/ |
|  | 158.61 ms  (66 slices for TR of 3.5s) | 95.71 ms  (108 slices for TR of 3.5s) |

**S3 Table**. **Performance of TR-external EPIK in comparison to EPI.** Under the same TE (22 ms) condition, the maximum matrix size achieved with EPI was 282 × 282, which is significantly smaller than that from TR-external EPIK (408 × 408). For the same matrix size (408 × 408), the minimum TE and TR required by EPIK was considerably smaller than that of EPI. This leads to a larger number of slices (i.e. larger brain coverage) in TR-external EPIK; that is, 42 slices more than that achievable with EPI when the TR is given as 3.5 s. The results above were obtained while other parameters were kept identical: FOV of 210 × 210 mm^2^, partial Fourier of 5/8 and parallel imaging of three-fold.

In EPI-based sequences, signal decay experienced during the relatively long readout induces image blurring and a concomitant deterioration of the desired spatial resolution, as defined by the image matrix size. Here, a quantitative evaluation of the spatial resolution was carried out when the above half-millimetre protocol was employed for both EPI and TR-external EPIK. S6 Figa shows the signal decay trajectories from the two techniques, simulated as a function of the k-space encoding index. For comparison purposes, the same TE (37 ms) – which is the minimum TE required for EPI (S3 Table) – was employed for both techniques. The T_2_^*^ value used in this simulation was 33.2 ms, which is known to be the T_2_^*^ of the grey matter at 7T [1]. The figure shows that the behaviour of signal decay for the central k-space region (i.e. keyhole) is the same for both techniques. However, due to the sparse sampling in EPIK, the signal decay for the peripheral k-space is much smaller in TR-external EPIK than in EPI. The point spread function from each signal decay trajectory was obtained as shown in S6 Figb. This plot shows that the PSF from TR-external EPIK is narrower than that of EPI. The full-width at half-maximum (FWHM) of the PSF was measured as 0.0174 and 0.0107 for EPI and TR-external EPIK, respectively; that is, an improvement of 39% for TR-external EPIK. The result infers that the *actual* spatial resolution of the nominal half-millimetre protocol is better when measured by TR-external EPIK than by EPI.

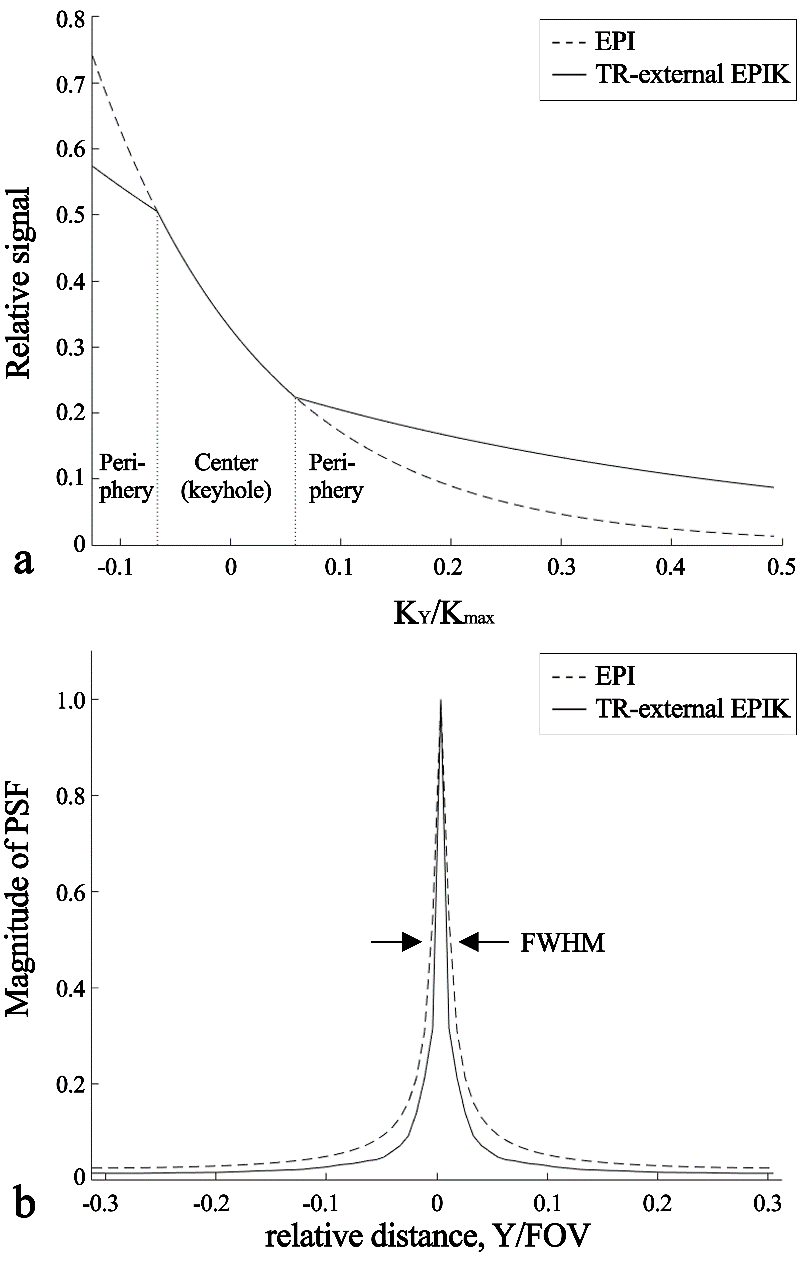

**S6 Fig. Simulation and valuation of spatial resolution for matrix size of 408 × 408.** (**a**) Signal decay trajectories with respect to the sampled k-space encoding index and (**b**) the resultant PSF associated with the signal decay trajectories in (a). The accumulated signal decay is less in TR-external EPIK, and accordingly, the PSF is narrower than that for EPI. The full-width at half-maximum (FWHM) of the PSF was measured to be 0.0174 and 0.0107 for EPI and TR-external EPIK, respectively.

In the main text, the imaging performance of TR-external EPIK was compared with several previous submillimetre-resolution fMRI studies in terms of brain coverage, single voxel volume and the number of voxels per temporal volume. S4 Table shows more details of the imaging parameters employed in each fMRI study. The improvements in TR-external EPIK against other fMRI studies can also be clearly seen here. The table additionally lists the base imaging method, TE and TR of each case. In particular, the maximum number of slices provided by each method can change depending on other imaging parameters, such as the employed matrix size, TE and TR. Since these parameters are not identical for all the works listed here, the performance of these methods in terms of slice throughput was approximated by computing the ‘TR/slice’ (the 6th column); the TR (i.e. for all slices) is also shown. This value was 32.41 ms for TR-external EPIK, which is slightly larger than the smallest case (26.19 ms; Kay^2019^). However, the larger ‘TR/slice’ in TR-external EPIK can be readily seen as being due to the in-plane matrix size of TR-external EPIK (408 × 408), which is significantly bigger than that employed by Kay^2019^ (200 × 162). Consequently, the increase in the number of voxels was 413.78% in TR-external EPIK, whilst the increase of the ‘TR/slice’ was only 23.75 % when compared to the previous work. The TE employed in each fMRI study was slightly different, ranging from 20 ms to 32 ms. The TE used here was 22 ms, which is nearly identical to the 2D EPI cases (Koizumi^2019^; Kay^2019^); TR-external EPIK is also based on a 2D echo-planar readout, although the sampling strategy is EPIK.

|  | **Imaging**  **method** | **FOV (mm^3^)**  **(Brain coverage (cm^3^))** | **Matrix size**  **(Number of Voxels)** | **Resolution (mm^3^)**  **(Voxel volume (mm^3^))** | **TR (ms)**  **(TR/slice)** | **TE** |
| --- | --- | --- | --- | --- | --- | --- |
| Heidemann^2012^ | Zoomed-GRAPPA | 109 × 156 × 19.5  (0331.58) | 169 × 240 × 30  (01.22 M) | 0.65 × 0.65 × 0.65  (0.27) | 3500  (116.67) | 27 |
| Kok^2016^ | 3D EPI | 192 × 192 × 38.4  (1415.58) | 240 × 240 × 48  (02.76 M) | 0.80 × 0.80 × 0.80  (0.51) | 3408  (071.00) | 28 |
| Kemper^2018^ | Inner-volume  3D-GRASE | 22.4 × 105 × 8.4  (0019.76) | 32 × 150 × 12  (00.06 M) | 0.70 × 0.70 × 0.70  (0.34) | 2000  (166.67) | 30.85 |
| Kasper^2019^ | 2D spiral | 220 × 220 × 36  (1742.40) | 275 × 275 × 36  (02.72 M) | 0.80 × 0.80 × 0.90  (0.58) | 3300  (091.67) | 20 |
| Kay^2019^ | 2D EPI | 160 × 129.6 × 67.2  (1393.46) | 200 × 162 × 84  (02.72 M) | 0.80 × 0.80 × 0.80  (0.51) | 2200  (026.19) | 22.4 |
| Koizumi^2019^ | 2D EPI | 123 × 188 × 32  (0739.97) | 154 × 236 × 40  (01.45 M) | 0.80 × 0.80 × 0.80  (0.51) | 2500  (062.50) | 21.8 |
| Sharoh^2019^ | 3D EPI | 215 × 215 × 100.8  (4659.48) | 228 × 228 × 112  (05.82 M) | 0.94 × 0.94 × 0.90  (0.80) | 3960  (035.36) | 20 |
| Dijk^2020^ | 3D multi-shot EPI | 131 × 120 × 24  (0377.28) | 187 × 171 × 84  (01.09 M) | 0.70 × 0.70 × 0.70  (0.34) | 4000  (047.62) | 28 |
| Huber^2020^ | VASO | 127 × 127 × 23.76  (0383.23) | 162 × 162 × 24  (00.63 M) | 0.79 × 0.79 × 0.99  (0.62) | 4400  (183.33) | 32 |
| Guidi^2020^ | VASO | 105 × 35 × 18  (0066.15) | 132 × 44 × 10  (00.06 M) | 0.80 × 0.80 × 1.80  (1.15) | 1648  (164.80) | 24 |
| **Proposed method** | 2D TR-external EPIK | 210 × 210 × 108  (4762.80) | 408 × 408 × 108  (17.98 M) | 0.51 × 0.51 × 1.00  (0.26) | 3500  (032.41) | 22 |

**S4 Table. Imaging configuration employed in several submillimetre-resolution fMRI studies.** This table shows the fMRI parameters employed in previous (2nd - 10th row) studies and the current work (the last row). All of the fMRI studies human brain studies performed at 7T. TR-external EPIK outperforms other methods in terms of brain coverage (4762.80 cm^3^), in-plane resolution (408 × 408; 0.51 × 0.51 mm^2^) and the number of voxels per temporal volume (17.98 M).
